# Deep Local Analysis deconstructs protein - protein interfaces and accurately estimates binding affinity changes upon mutation

**DOI:** 10.1101/2022.12.04.519031

**Authors:** Yasser Mohseni Behbahani, Elodie Laine, Alessandra Carbone

**Affiliations:** Sorbonne Université, CNRS, IBPS, Laboratory of Computational and Quantitative Biology (LCQB), UMR 72308, Paris, 75005, France

## Abstract

The spectacular recent advances in protein and protein complex structure prediction hold promise for reconstructing interactomes at large scale and residue resolution. Beyond determining the 3D arrangement of interacting partners, modeling approaches should be able to unravel the impact of sequence variations on the strength of the association. In this work, we report on Deep Local Analysis (DLA), a novel and efficient deep learning framework that relies on a strikingly simple deconstruction of protein interfaces into small locally oriented residue-centered cubes and on 3D convolutions recognizing patterns within cubes. Merely based on the two cubes associated with the wild-type and the mutant residues, DLA accurately estimates the binding affinity change for the associated complexes. It achieves a Pearson correlation coefficient of 0.81 on more than 2 000 mutations, and its generalization capability to unseen complexes is higher than the state-of-the-art methods. We show that taking into account the evolutionary constraints on residues contributes to predictions. We also discuss the influence of conformational variability on performance. Beyond the predictive power on the effects of mutations, DLA is a general framework for transferring the knowledge gained from the available non-redundant set of complex protein structures to various tasks. For instance, given a single partially masked cube, it recovers the identity and physico-chemical class of the central residue. Given an ensemble of cubes representing an interface, it predicts the function of the complex. Source code and models are available at http://gitlab.lcqb.upmc.fr/DLA/DLA.git.

## 1 Introduction

The ever-growing number of sequenced individual genomes and the possibility of obtaining high-resolution 3D structural coverage of the corresponding proteomes [46, 36] opens up exciting avenues for personalized medicine. Assessing the impact of sequence variations, particularly missense mutations, between individuals on how proteins interact with each other can shed light on disease susceptibility and severity [51, 35, 68, 10] and help decipher gene-disease-drug associations for developing therapeutic treatments [24, 49, 65, 55, 30, 67]. Of particular interest are the surface regions of proteins directly involved in the interactions, as this is where most disease-related missense mutations occur. [74, 13, 12, 22, 33]. At the same time, rapid advances in deep learning techniques for biology, especially for biomolecules, are creating opportunities to revisit the way we look at protein complexes and represent them.

The impact of a mutation on the strength of the association between two protein partners can be measured by the difference in binding free energy

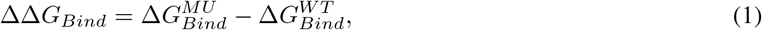

where 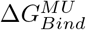 and 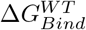 are the binding free energies, or binding affinities, of the mutated and wild-type complexes, respectively. Significant efforts have been expended over the past decade to produce, collect and curate binding affinity measurements for wild-type and mutated complexes (**Table S 1)** [31, 41, 63, 32, 72, 47]. Nevertheless, the handful of experimental techniques yielding accurate estimates of Δ*G_Bind_* remain laborious, expensive and time-consuming [70]. To overcome this limitation, several efficient computational methods have been developed (**Table S 2)** [60, 78, 76, 73, 42, 20, 59, 3, 75, 54, 23]. Most of them exploit local environments around the mutation site to directly predict ΔΔ*G_Bind_* values. The advantage of this strategy is twofold. First, it avoids the accumulation of errors on the 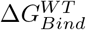 and 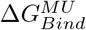 quantities that would result in large approximations in ΔΔ*G_Bind_*. Second, it avoids the unnecessary calculation of properties not modified by the mutation, *e.g.,* the chemical composition of the non-interacting surface and the 3D geometry of the interface contact distribution. Indeed, these properties, while contributing strongly to the binding affinity [57, 70], are not, or only slightly, sensitive to point mutations located at the interface. The state-of-the-art methods sometimes achieve very high prediction accuracy, but their ability to generalize to diverse complexes can be improved [20].

Representation learning powered by deep neural networks has opened up extraordinary opportunities to develop all-purpose models transferring knowledge across systems and tasks. After a major breakthrough in natural language processing [14, 71], the concept has been transferred to proteins through protein language models (pLMs) [58, 17, 4, 26]. pLMs learn the fundamental properties and mechanisms of natural protein diversity by reconstructing some masked or the next amino acid(s), given their sequence context, at scale. They exhibit exciting potential for a broad range of protein-related problems [40, 62, 43, 44, 56, 78]. Beyond sequence information, self-supervised learning-based approaches have leveraged the protein and protein complex 3D structures available in the Protein Data Bank (PDB) [5] for fixed-backbone protein design [2, 28, 11], for predicting protein stability [6, 77], and for assessing the impact of mutations on protein-protein interactions [42]. In particular, in [42], a graph neural network is trained to reconstruct disturbed wild-type and mutated complex structures represented as graphs. A gradient-boosting trees algorithm then exploits the learned representations to predict mutation-induced ΔΔ*G_Bind_* values. Although this approach showed promising results, it sequentially employs two different machine learning components trained independently, limiting its versatility and applicability to other tasks.

Here, we report on *Deep Local Analysis*(*DLA*)-*Mutation*, the first end-to-end deep learning architecture estimating mutation-induced ΔΔ*G_Bind_* from patterns in local interfacial 3D environments learnt through self-supervision (**Fig. 1)**. It builds on the DLA framework we previously introduced for assessing the quality of protein complex conformations [48]. DLA applies 3D convolutions to locally oriented residue-centred cubes encapsulating atomic-resolution geometrical and physico-chemical information [52] (**Fig. 1A**). In this work, we expanded this framework by combining self-supervised representation learning of 3D local interfacial environments (**Fig. 1B**) with supervised learning of ΔΔ*G_bind_* exploiting both structural and evolutionary information (**Fig. 1C**). DLA-Mutation only takes as input two cubes, corresponding to the environments around the wild-type and mutated residues, respectively, and directly estimates ΔΔ*G_bind_.* Beyond prediction, we used the learned representations to investigate the extent to which the environment of an interfacial residue is specific to its type and physico-chemical properties (**Fig. 1D**). DLA-mutation code and models are freely available to the community at http://gitlab.lcqb.upmc.fr/DLA/DLA.git.

**Figure 1:**
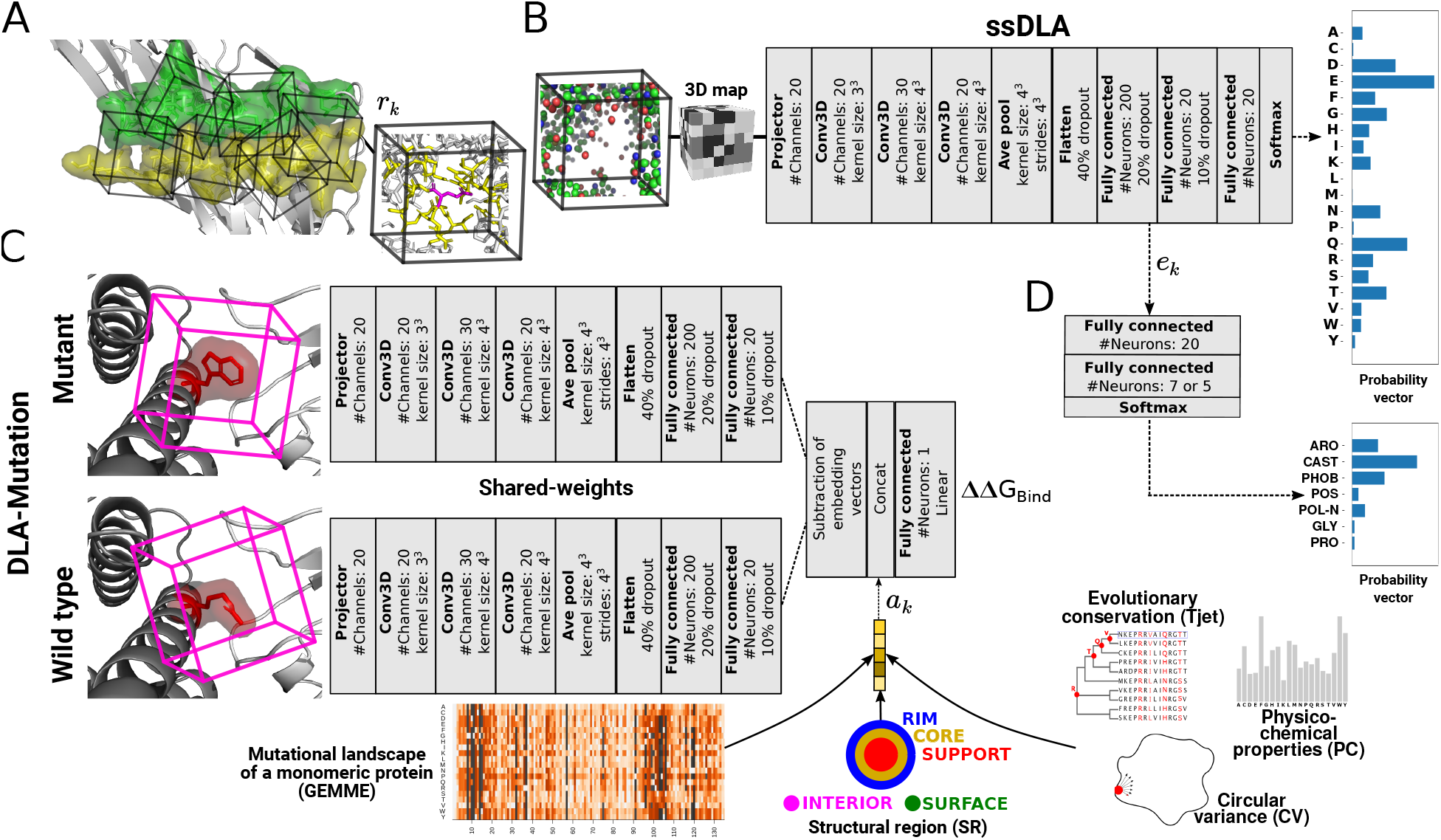
DLA data representations and architectures. **A.** A representation of a protein interface (green and yellow residues from each partner) as an ensemble of cubes (*I_C_*). Each cube (*r_k_* ∈ *I_C_*) is centred and oriented around an interfacial residue. In the example cube on the right, the atoms displayed in yellow and magenta sticks are enclosed in a 5 Å-radius sphere centred on a randomly chosen atom from the central residue. **B.** Architecture of the self-supervised model, named ssDLA. The input cube is the same as in panel A. The atoms that were in transparent sticks are now replaced by an empty space. Carbon atoms are colored in green, oxygen in red, nitrogen in blue, and sulfur in yellow. The training task is to recover the identity of the residue lying at the center of the partially masked input cube. **C.** Siamese architecture of the supervised model DLA-Mutation predicting mutation-induced binding affinity changes. The two parallel branches with shared weights apply 3D convolutions to the local 3D environments around the wild-type and mutated residues and compute two embedding vectors. Auxiliary features are concatenated to the vector resulting from subtracting these two embedding vectors. **D.** Two-layer dense classifier taking as input the embedding vectors computed by the pre-trained ssDLA (panel B) and outputting a probability vector whose dimension is the number of classes.

## 2 Methods

### 2.1 Protein–protein interface representation

We represent a protein-protein interface as a set of locally oriented cubic volumetric maps centered around each interfacial residue (**Fig. 1A**). The local atomic coordinates of the input structure are first transformed to a density function, where each atom is one-hot encoded in a vector of 167 dimensions [52]. Then, the density is projected on a 3D grid comprising 24 × 24 × 24 voxels of side 0.8Å. The map is oriented by defining a local frame based on the common chemical scaffold of amino acid residues in proteins [52] (see Supplementary Information for more details). This representation is invariant to the global orientation of the structure while preserving information about the atoms and residues relative orientations.

For the self-supervised representation learning, we trained DLA to recognize which amino acid would fit in a given local 3D environment extracted from a protein-protein interface. Our aim in doing so is to capture intrinsic patterns underlying the atomic arrangements found in local interfacial regions. Formally, the machine predicts the probability *P*(*y*|*env*) of the amino acid type *y*, for *y* ∈ {*A, C, D,…,W,Y*}, conditioned on the interfacial local chemical environment *env* given as input. In practice, we process the input cube before giving it to DLA by masking a sphere of radius r_c_Å centered on an atom from the central residue (**Fig. S 4** and **Fig. 1A**). Masking a fixed volume prevents introducing amino acid-specific shape or size biases. We experimented with different values of *r_c_* (3 and 5Å) and different choices for the atom (*C_α_, C_β_*, random). We found that a sphere of radius of 5Å with a randomly chosen center yielded both good performance and expressive embedding vectors.

For the supervised prediction of ΔΔ*G_bind_*, we combined the embedding vectors of the volumetric maps with five pre-computed auxiliary features (**Fig. 1C**), among which four describe the wild-type residue, namely its conservation level *T_jet_* determined by the Joint Evolutionary Trees method [18], its physico-chemical properties to be found at interfaces (*PC*), its protruding character, as measured by its circular variance (*CV*) [45, 7], and the structural region (SR) where it is located: protein interior (INT), the non-interacting surface (SUR), or, if it is part of the interface, the support (S or SUP), the core (C or COR), or the rim (R or RIM) as defined in [39]. We previously demonstrated the usefulness of these properties for predicting and analysing protein interfaces with other macromolecules (protein, DNA/RNA) [48, 9, 57, 37]. The fifth feature is a numerical score computed by GEMME [38] that reflects the impact of the point mutation on the function of the protein chain where it occurs, considered as a monomer. GEMME combines the conservation levels *T_jet_* with amino acid frequencies and the minimum evolutionary distance between the protein sequence and an homologous protein displaying the mutation. See Supplementary Information for more details.

### 2.2 DLA architectures

Our DLA framework is relatively simple, generic and versatile. Its main core architecture comprises a projector, three 3D convolutional layers, an average pooling layer, and a fully connected subnetwork (**Fig. 1B-C**). The purpose of the projector is to reduce the dimension of each input cube voxel’s feature vector from 167 to 20. We apply batch normalization after each 3D convolution. The average pooling layer exploits scale separability by preserving essential information of the input during coarsening of the underlying grid. To avoid overfitting, we applied 40%, 20%, and 10% dropout regularization to the input, the first and the second layers, respectively, of the fully connected subnetwork.

For the self-supervised task (**Fig. 1B**), the fully connected subnetwork contains three successive layers (sizes 200, 20 and 20) and the last activation function (Softmax) outputs a probability vector of size 20 representing the 20 amino acids. The categorical cross-entropy loss function measures the difference between the probability distribution of the predicted output and a one-hot vector encoding the true amino acid type of the central residue. We refer to this version of DLA to build the pre-trained model as *self supervised-DLA* or *ssDLA.*

For the supervised ΔΔ*G_bind_* prediction (**Fig. 1C**), we used the core DLA framework to build a Siamese architecture constituted by two branches with shared weights. The network processes two input cubes corresponding to the wildtype and mutated residues. The average pooling layer is followed by two fully connected layers of size 200 and 20, respectively, within each branch. We then merge the two branches by subtracting the computed embedding vector and concatenate the auxiliary features (described above) to the resulting vector. The last fully-connected layer displays a linear activation function and outputs one value. The loss is the mean squared error. We refer to this architecture as *DLA-Mutation.*

### 2.3 Databases

We computed the ground-truth ΔΔ*G_bind_* values from SKEMPI v2.0 [31], the most complete source for experimentally measured binding affinities of wild-type and mutated protein complexes. We restricted our experiments to the data produced by the most reliable experimental techniques, namely Isothermal Titration Calorimetry (ITC), Surface Plasmon Resonance (SPR), Spectroscopy (SP), Fluorescence (FL), and Stopped-Flow Fluorimetry (SF), as done in [70]. We selected a subset of 2 003 mutations associated with 142 complexes, referred to as *S2003* in the following. To provide ssDLA and DLA-Mutation with input protein-protein complex 3D structures, we created and processed two databases, namely *PDBInter* and *S2003-3D.* PDBInter is a non-redundant set of 5 055 experimental structures curated from the PDB. S2003-3D contains 3D models generated using the “backrub” protocol implemented in Rosetta [64]. We refer to each generated conformation as a backrub model. We generated 30 backrub models for each wild-type or mutated complex. This amount was shown to be sufficient for estimating free energies in [3]. See Supplementary Information for more details.

### 2.4 Training and evaluation of ssDLA and DLA-Mutation

We trained and validated ssDLA on the PDBInter database. The protein complexes in the train set do not share any family level similarity with the 142 complexes from S2003, according to the SCOPe hierarchy. We generated 247 662 input samples (interfacial cubes) from the train set and 34 174 from the validation set. Amino acids are not equally distributed in these sets; leucine is the most frequent one, while cysteine is the rarest (**Fig. S 3)**. To compensate for such imbalance and with the aim of penalizing more those errors that are made for the less frequent amino acids, we assigned a weight to the loss of each amino acid type that is inversely proportional to its frequency of occurrence (**Table S 4)**. We trained ssDLA for 50 epochs with the Adam optimiser in TensorFlow [1] at a learning rate of 0.0001 (**Fig. S 5A**). We explored different hyperparameter values by varying the learning rate, applying different normalisation schemes, changing the compensation weights, etc. We retained the hyperparameters leading to the best performance on the validation set. The trained ssDLA model extracts *embedding vector e_k_* of size 200 (**Fig. 1B**) for a given cube.

We used S2003 to train and test DLA-Mutation. We set the learning rate at 0.001 and we initialized the weights of the network with those of the pre-trained ssDLA model. We first evaluated DLA-Mutation through a 10-fold cross validation performed at the mutation level. This evaluation procedure, which is widely used in the literature [20, 59, 60, 73, 78], considers each sample independently when splitting the data between train and test sets (*mutation-based* split). However, this assumption is problematic since the same complex or even the same wild-type residue may be seen during both the training and the testing phases. These cases are expected to be “easy” to deal with. For a more challenging and realistic assessment, we held out 32 complexes displaying 391 mutations for the testing phase, and trained DLA-Mutation on the rest of the dataset (*complex-based* split).

For the comparison with iSEE, we used the same train and test procedure as that reported in [20] (**Table S 2)**, using the wild-type and mutant 3D models produced by HADDOCK [69] and available from [20]. For the comparison with the other predictors, we defined the test set from the intersection between S2003 and the benchmark set used in [20]. It amounted to 112 mutations from 17 complexes. We defined a new training set comprising 945 mutations from S2003 coming from complexes sharing less than 30% sequence identity with those from this test set. In the case of GraphPPI [42] and TopNetTree [73], the comparison remains qualitative due to the lack of software package from GraphPPI and the function-specific model for TopNetTree that is trained only on antibody-antigen complexes from the AB-Bind database [63].

### 2.5 Mapping the embeddings to residue and interface properties

We trained a fully-connected network comprised of only 1 hidden layer of size 20 to map the embeddings computed by ssDLA to residue- and interface-based properties. The input layer is of size 200 and the Softmax activation function of the output layer computes a probability vector whose size is the number of classes. We used categorical cross-entropy as the loss function. In the first experiment we mapped an input embedding vector (*e_k_*, size 200, see **Fig. 1D**), representing a local 3D interfacial environment, to an output amino acid physico-chemical class, among the seven defined in [38] (**Table S 3)**. We directly gave the embedding vector computed by ssDLA for a given input cube to the classifier. In the second experiment, we mapped an input embedding vector averaged over an entire interface to an output interaction functional class, among antibody-antigen (AB/AG), protease-inhibitor (Pr/PI) and T-cell receptor - major histocompatibility complex (TCR/pMHC), as annotated in the SKEMPI v2.0 database. For training purposes, we redundancy-reduced the set of 142 complexes from *S2003* based on a 30% sequence identity cutoff. See Supplementary Information for details.

## 3 Results

The DLA framework deconstructs a protein-protein interface to predict mutation-induced changes in binding affinity and solve residue- or interface-based downstream tasks (**Fig. 1)**. It extracts embedding vectors from locally oriented cubes surrounding wild-type or mutant interfacial residues and combines them with auxiliary features, including structural regions or evolutionary information.

### 3.1 DLA-Mutation accurately predicts ΔΔ*G_bind_*

DLA-Mutation achieved an overall very good agreement with ΔΔ*G_bind_* experimental measurements (**Fig. 2)**. It reached a Pearson correlation coefficient (PCC) of 0.812 over 2 003 single-point mutations (**Fig. 2A**), following a mutation-based 10-fold cross validation (see *Methods*). Moreover, it showed high generalization capability to unseen complexes, achieving a PCC of 0.735 and a root mean squared error (RMSE) of 1.23 kcal/mol (**Fig. 2B** and **Table 1)**, following a complex-based train and test split procedure (see *Methods*).

**Figure 2:**
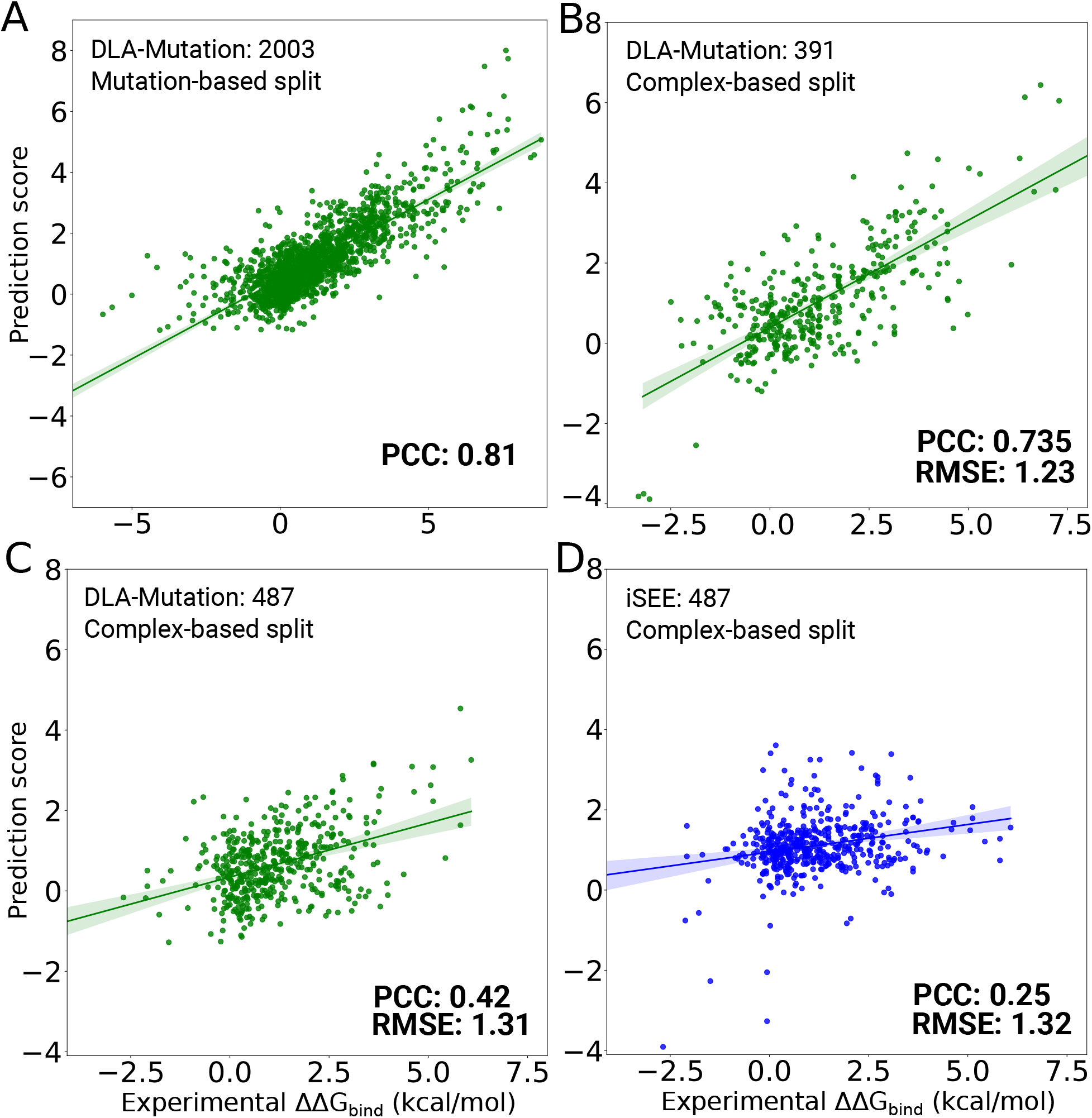
Performance of DLA-Mutation and comparison with iSEE. **A-B.** DLA-Mutation scores versus experimental ΔΔ*G_bind_* values on S2003 dataset. **A.** Mutation-based 10-fold cross validation procedure over all 2003 mutations (see *Methods*). The model relies on fine-tuned weights, starting from those of the pre-trained ssDLA, and exploits all auxiliary features. **B.** Test set of 391 mutations coming from 32 complexes that were not seen during training and were randomly selected from the S2003 dataset. **C-D.** DLA-Mutation (green) and iSEE (blue) scores versus experimental ΔΔ*G_bind_* values for the test set of 487 mutations from 56 complexes (S487 dataset). The input 3D models and training and evaluation procedure were directly taken from [20]. **C.** DLA-Mutation **D.** iSEE.

**Table 1:** Different experimental setups for ΔΔ*G_bind_* prediction with DLA-Mutation.

| Train/Test split level | #(mutations) | | Weight initialization | Auxiliary features | PCC | RMSE ($\frac{kcal}{mol}$) |
| --- | --- | --- | --- | --- | --- | --- |
|  | Train | Test |  |  |  |  |
| mutation | 2003 | - | pre-training | *All* | 0.812 | - |
| complex | 1612 | 391 | pre-training | *SR* | 0.686 | 1.31 |
|  |  |  |  | *SR-Tjet* | 0.712 | 1.32 |
|  |  |  |  | *SR-GEMME* | 0.726 | 1.27 |
|  |  |  |  | *All* | 0.735 | 1.23 |
|  |  |  | random | *SR* | 0.602 | 1.44 |
|  |  |  |  | *All* | 0.657 | 1.37 |
| complex and <30% seq. id. | 945 | 112 | pre-training | *All* | 0.481 | 1.14 |
| complex | 1102* | 487* | pre-training | *All* | 0.423 | 1.31 |

\* The input 3D models, generated by HADDOCK [69], were taken from [20].

### 3.2 Evolutionary information and pre-training matter

To build the DLA-Mutation model, we fine-tuned the weights of the pre-trained ssDLA model (**Fig. 1B**) to predict ΔΔ*G_bind_* values in a supervised fashion (**Fig. 1C**). For each mutation, we combined information coming from the local 3D environments of the wild-type and mutant residues extracted from 3D models of the corresponding complexes (S2003-3D database, see *Methods* and **Fig. S 1)** with information about the structural region of the wild-type residue (**Fig. 1C**, *SR*), as well as other geometrical (*CV*), physico-chemical (*PC*), and evolutionary (*GEMME, T_jet_*) descriptors. We performed an ablation study to assess the contribution of these descriptors and of the pre-training step (**Fig. S 7** and **Table 1)**.

In the baseline configuration, the only auxiliary feature we used was *SR*. We have previously shown that this information contributes significantly to the performance of the DLA framework [48]. In addition to this, we also considered evolutionary information, using either *GEMME* scores (*SR-GEMME*) or *T_jet_* conservation levels (*SR-Tjet*). We found that the wild-type residue’s buriedness (*CV*) and interface propensity (*PC*) contributed very little to the accuracy of the predictions (**Fig. S 7A-D** and **Table 1,** compare *All* with *SR-T_jet_* and *SR-GEMME*). Removing them is essentially harmless. By contrast, evolutionary information does significantly contribute to the model’s performance, as attested by the rather low PCC (0.648) obtained when using only SR (**Fig. S 7B** and **Table 1)**. By design, the mutation-specific *GEMME* score is correlated to the position-specific conservation level *T_jet_* [38], and thus the two descriptors are redundant to some extent. Nevertheless, we observed that the former was more informative than the latter (**Fig. S 7,** compare panels C and D). Finally, pre-training the architecture through self-supervision with ssDLA clearly improved the predictions (**Fig. S 7,** compare panels A-B with panels E-F, and **Table 1)**. The gain in PCC is of 0.08 compared to initialising DLA-Mutation weights randomly.

### 3.3 Comparison with state-of-the-art predictors

We further assessed DLA-Mutation generalisation capabilities with respect to iSEE [20], a recently developed machine learning-based method (**Fig. 2C-D**). Similarly to DLA-Mutation, iSEE directly estimates ΔΔ*G_Bind_* values exploiting structural information coming from the wild-type and mutant complex 3D structures, as well as evolutionary information. We found that DLA-Mutation generalised better than iSEE from SKEMPI version 1 to version 2 (**Fig. 2C-D**). Specifically, when trained on SKEMPI v1.0, it reached a PCC of 0.423 on 487 mutations coming from 56 unseen complexes from SKEMPI v2.0 (**Fig. 2C**). The correlation obtained with iSEE was much lower, around 0.25 (**Fig. 2D**). In this experiment, the train and test conditions were exactly the same for the two methods, with the same input HADDOCK-generated 3D models [20]. We then extended the comparison to three other ΔΔ*G_bind_* predictors, namely mCSM [53], FoldX [23] and BindProfX [75] (**Fig. 3)**. mCSM directly estimates ΔΔ*G_Bind_* values by exploiting the 3D structure of the wild-type complex and descriptors of the substituting amino acid within a machine learning framework. FoldX estimates free energies of binding Δ*G_Bind_* of the wild-type and mutant complexes using a physics-based energy function and then computes their difference. BindProfX combines FoldX with evolutionary interface profiles built from structural homologs. DLA-Mutation outperforms all of the predictors on a set of 112 mutations coming from 17 complexes sharing less than 30% sequence identity with those seen during training (**Fig. 3)**. Finally, we considered two recent deep learning-based approaches, namely GraphPPI [42] and TopNetTree [73]. The reported performance for these tools are similar to those we obtained for DLA-Mutation (**Table S 2)**. They both experience a drop in performance from the mutation-based cross validation to the blind test on unseen complexes. Contrary to DLA-Mutation, they do not predict ΔΔ*G_Bind_* in an end-to-end fashion. Moreover, they take the full complex structure as input, instead of local environments.

**Figure 3:**
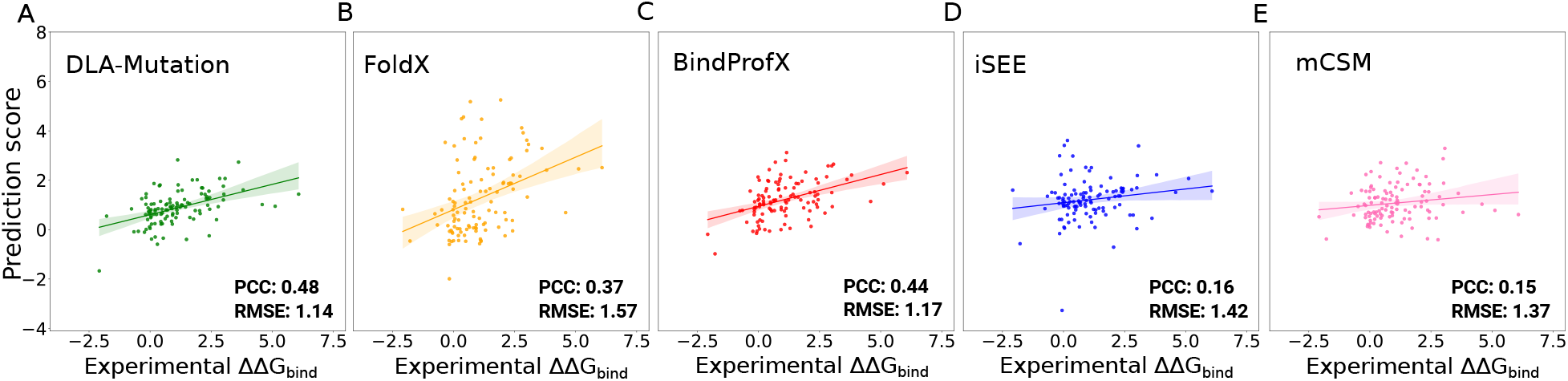
Comparison between DLA-Mutation and other ΔΔ*G_Bind_* predictors. We report values for 112 mutations coming from 17 protein complexes not seen during the training or optimisation of any of the predictors. **A.** DLA-Mutation was trained on 945 mutations from S2003 coming from complexes sharing less than 30% sequence identity with those from this test set. We used fine-tuning of the weights and all auxiliary features. **B-E.** The scores reported for FoldX (B), BindProfX (C), iSEE (D) and mCSM (E) were taken directly from [20].

### 3.4 DLA-Mutation performs better on core and rim residues and is robust to size and sequence identity changes

The location of a mutation in a protein interface might be a relevant indicator for the confidence in the estimation. We investigated this issue by describing an interface as three concentric layers of residues, the support (internal layer), the core (the second layer) and the rim (the third and most external layer) [39]. DLA-Mutation better deals with mutations taking place in the core and rim, with PCCs as high as 0.737 and 0.798, respectively (**Fig. 4A**, compare gold and blue dots with red dots). The mutations in the core are also the most frequent ones. By contrast, very few mutations are located outside of the interface and the associated range of experimental ΔΔ*G_Bind_* values is very narrow, making it difficult to distinguish them (**Fig. 4A**, pink and green dots). In addition, the prediction accuracy seems to depend on the function of the complex, with the protease-inhibitor class displaying the highest number of complexes and the highest accuracy (**Fig. 4B**). However, this observation may be interpreted in the light of the nature of the substitutions. Indeed, the protease-inhibitor complexes display a wide variety of substitutions, while the other classes mostly display substitutions to alanine. Overall the predictions are more accurate when the mutant amino acid is not alanine (PCC of 0.790 versus 0.34). The amino acid size change itself is not a determining factor (**Fig. 4C**). DLA-Mutation performs consistently well on small-to-large, large-to-small and size-neutral substitutions, with a slight preference for the latter (PCC = 0.793). Finally, the predictions are robust to variations in the sequence identity between the test and train complexes (**Fig. 4D**).

**Figure 4:**
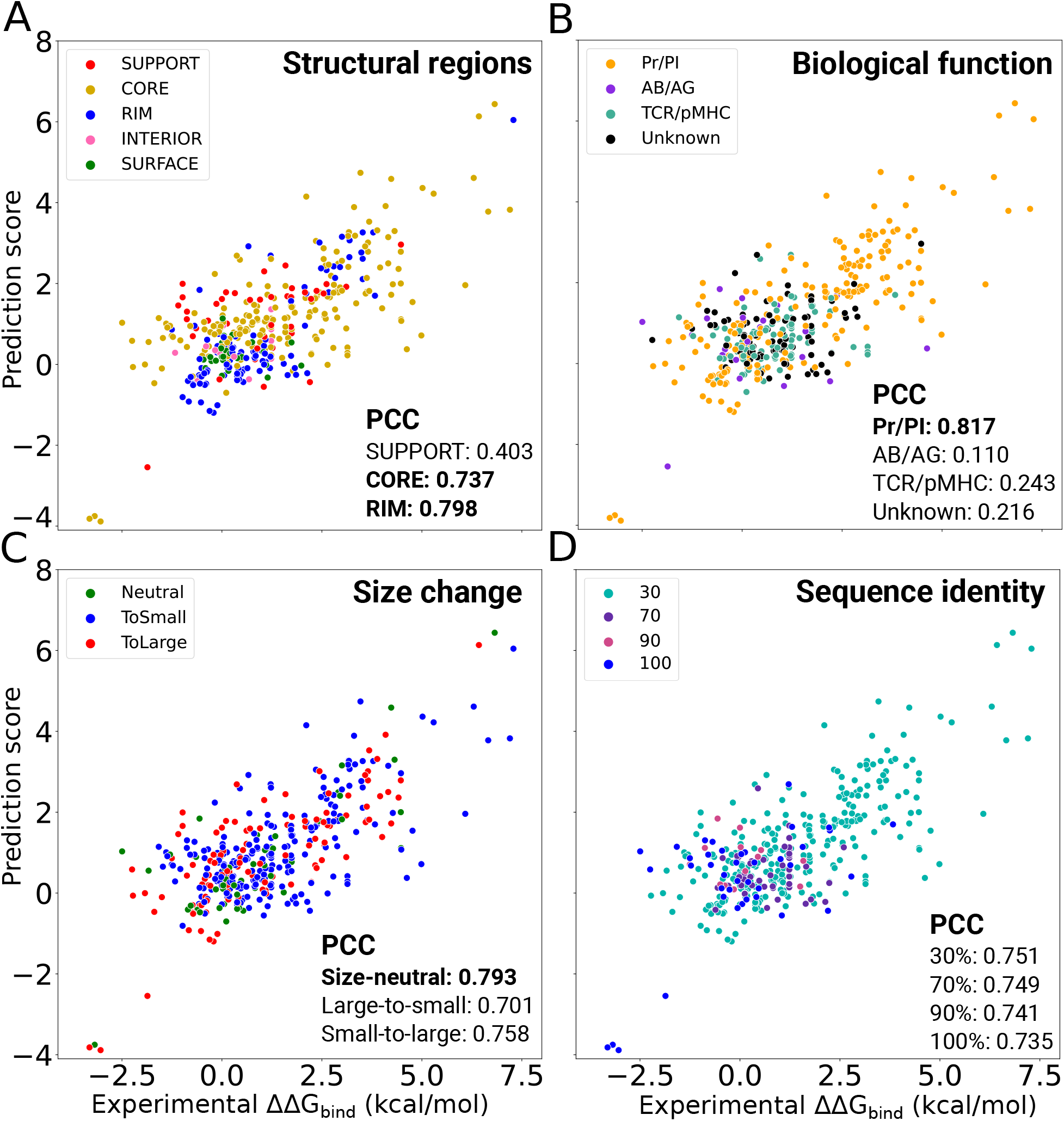
Influence of different residue-based and complex-based properties on DLA-Mutation accuracy. The predicted and experimental values are reported for 391 mutations coming from 32 complexes not seen during training (randomly selected from S2003 dataset). We used weight fine-tuning and all auxiliary features. The overall PCC is 0.720 (**Table 1)**. The dots are colored with respect to the structural region where the mutated residue lies (**A**), the complex’s biological function (**B**), the amino acid size change upon mutation (**C**) and the minimum sequence identity shared with any training complex (**D**). We calculated the change of amino acid size as a volume difference (*δV*) between wild-type and mutant following [25]. A mutation was classified as size-neutral if |*δV*| < 10Å, as small-to-large if *δV* > 10Å, and as large-to-small if *δV* < −10Å.

### 3.5 Can an interfacial residue be learnt from its environment?

ssDLA’s ability to recover the identity of the central residue in the input cube can inform us about the extent to which an interfacial residue’s 3D environment is specific to its amino acid type or physico-chemical properties. To investigate this possibility, we analysed the probability vectors computed by ssDLA when given a partially masked cube as input (**Fig. 5A**). To avoid any amino acid-specific bias, we masked a volume of constant shape and size, namely a sphere of radius 5Å, in all training samples (see *Methods* and **Fig. S 4)**. ssDLA successfully and consistently recognised the amino acids containing an aromatic ring (F, Y, W, H) and most of the charged and polar ones (E, K, R, and to a lesser extent Q and D), as well as methionine (M), cysteine (C), glycine (G), and proline (P), whatever their structural region (**Fig. 5A**). By contrast, the location of alanine (A), isoleucine (I) and leucine (L) influenced their detection. While they were ranked in the top 3 in the support and the core, they were almost never recognised in the rim. Inversely, the polar asparagine (N) was recognised when located in the rim or the core, but not the support. The model often confused the hydroxyl-containing serine (S) and threonine (T) on the one hand, and the hydrophobic I and L on the other hand.

**Figure 5:**
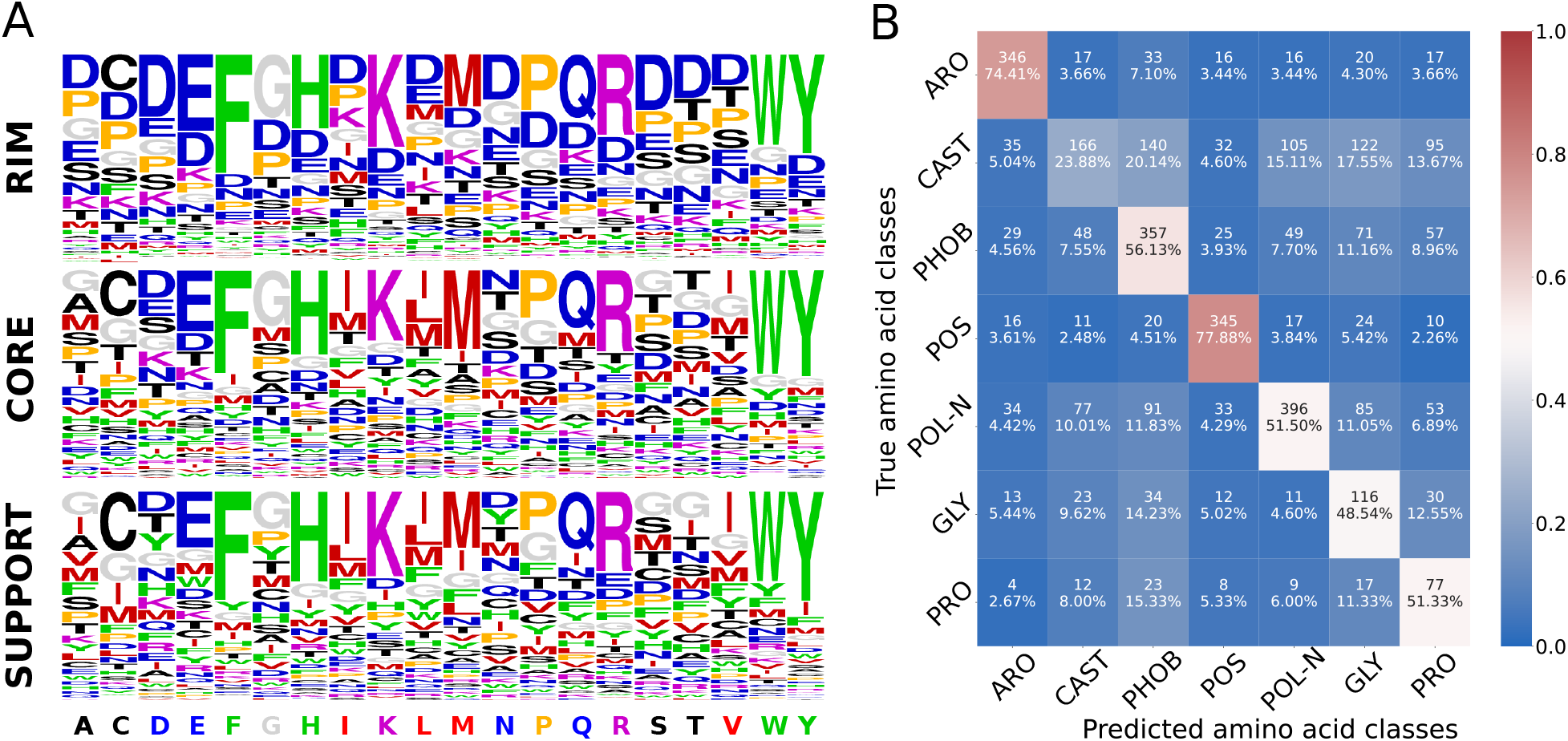
Performance of ssDLA model. **A.** The predictive power of ssDLA model is evaluated on the validation set of *PDBInter.* The three logos represent the propensities of each amino acid to be predicted (having maximum score in the output layer), depending on the true amino acid (x-axis) and on its structural region (see *Methods*. Amino acids are colored based on seven similarity classes: ARO (F, W, Y, H) in green, CAST (C, A, S, T) in black, PHOB (I, L, M, V) in red, POS (K, R) in purple, POL-N (N, Q, D, E) in blue, GLY (G) in gray and PRO (P) in orange (see *Methods*). **B.** Confusion matrix for the prediction of the 7 amino acid classes using embedding vectors generated by ssDLA. The percentage values and the colors indicate recall. The model is trained and tested on the interfacial residues of X-ray crystal structures of S2003. See **Fig. S 6** for the performance of ssDLA when only 4 channels corresponding to the four amino acid-independent chemical elements (O, C, N and S) are considered to define the cubic volumetric maps.

These tendencies cannot be deduced from the relative frequencies of occurrence of the different amino acids in the three interface structural regions (**Fig. S 3)**. For instance, ssDLA behaves very differently with N and Q (**Fig. 5A**), although they display the same relative abundances and the same structural regions preferences (**Fig. S 3)**. Hence, the poor recovery rate for N suggests that the environments for this amino acid are more ambiguous or diverse than those observed for Q. Likewise, ssDLA tendency to over-populate the rim with aspartate (D) does not reflect its overwhelming presence in this region. We hypothesise that D serves as a « bin » class predicted when the environment is underdetermined. Such underdetermination or ambiguity is more likely to happen in the rim, where the residues are more exposed and thus the cube contains more empty space. A previous study reported different trends for a similar task and similar data representation [2]. In particular, the model from [2] could identify G and P with very high success, whereas it confused F, Y and W. These results may reflect a bias toward recognising amino acid-specific sizes and shapes, due to masking only the side chain of the central residue. Moreover, the model was trained and evaluated on monomeric proteins.

### 3.6 Predicting residue- and interface-based properties

To evaluate the embedding vectors computed by the pre-trained ssDLA, we tested whether they could be mapped to per-residue and per-interaction physico-chemical and functional properties. To do so, we added a 2-layer fully connected network on top of ssDLA’s architecture (**Fig. 1D**), and we trained it to perform two downstream tasks (see *Methods*) The first task consisted in assigning amino acid physico-chemical classes to the input cubes. The amino acid classification we chose previously proved relevant for predicting the functional impact of mutations [38]. It distinguishes the aromatic amino acids (ARO: F, W, Y, H), the hydroxyl-containing ones plus alanine (CAST: C, A, S, T), the aliphatic hydrophobic ones (PHOB: I, L, M, V), the positively charged ones (POS: K, R), the polar and negatively charged ones (POL-N: N, Q, D, E), glycine (GLY), and proline (PRO) (**Table S 3)**. The per-class tendencies are consistent with those observed for the pre-training task (**Fig. 5,** compare the two panels). Specifically, the best performances are observed for the aromatic (ARO) and positively charged (POS) classes, with more than 70% recall, while the CAST class is the most difficult to identify. Conformational sampling influences the results. We observed improved performances when dealing with 3D models compared to experimental structures (**Fig. S 10)**. We may hypothesize that the backbone rearrangements and side-chain repacking performed by the backrub protocol lead to a better fit between the central amino acid and its environment (compare panels A and B). Averaging the embedding vectors over 30 models allows extracting with an even higher precision the intrinsic properties of the central amino acid (compare panels B and C). The second task was to predict the function of a protein-protein interaction. The embedding vectors proved useful to distinguish the protease-inhibitor assemblies (recall = 83.33%) from the two other functional classes (**Fig. S 11)**. The classifier tends to confuse the antibody-antigens with T-cell receptor-major histocompatibility complexes. This behaviour is expected, owing to the structural similarity shared between T-cell receptors and antibodies.

## 4 Discussion

Fundamental questions about protein-protein interactions require knowledge acquisition and transfer from protein-protein interfaces with deep learning approaches. In this work, we proposed a simple conceptual framework based on a small cube defined around an interfacial residue to deconstruct and decrypting protein-protein interactions. Our approach leverages the non-redundant set of experimentally resolved protein complex structures to assess the impact of mutations on protein-protein binding affinity, among other applications. It derives and contrasts representations of the local geometrical and physico-chemical environments around the mutation site in wild-type and mutated forms with a Siamese architecture. It complements the information embedded in these environments with evolutionary information from sequences related to the protein carrying the mutation. Compared to other state-of-the-art predictors, DLA-Mutation generalizes better to unseen complexes.

Despite the improvement over the state-of-the-art, the DLA-Mutation generalization capability from the first to the second version of SKEMPI remains limited. This result likely reflects differences in the protocols employed to produce, collect and manually curate the data between the older version, released in 2012, and the new one, released seven years later. More generally, ΔΔ*G_bind_* measurements may contain errors, *e.g.,* coming from systematic bias. Moreover, in SKEMPI v2.0, we observed that for some mutations, distinct values of mutant binding affinity were measured by different laboratories or using different experimental techniques [31].

Beyond predictive power, we have shown that the learned representations can help to better understand protein interfaces. Depending on the amino acid and the location at the interface, the 3D environment may be more or less ambiguous and diverse. We designed a constant volume masking procedure to avoid amino acid-specific size and shape biases. Nevertheless, the spherical mask of radius 5Å may not always cover the whole central residue, raising the question of whether the network relies on the amino acid-specific types of the remaining atoms in such cases. To test this, we removed any amino acid-specific information by reducing the 167 feature channels encoding the atom types to 4, corresponding to the four chemical elements C, N, O, and S. Even with four channels, ssDLA successfully recognized and distinguished the large aromatic amino acids F, W, and Y, as well as the long positively, charged R and K, whatever the structural region (**Fig. S 6)**. Besides reducing the number of channels, we also slightly lowered the weight of D in the calculation of the loss during training (**Table S 4)**. This small change shifted the tendency of ssDLA to predict D for E, especially in the rim region (**Fig. S 6)**. Such instability highlights the under-determination of the environments in this region. One direction of improvement for the ssDLA model would be to expand the set of training samples. Here, we limited the training to a “compacted” version of the ensemble of complex structures available in the PDB. Considering more structures could help the model learn residue-specific pattern variations and improve prediction performance.

The resolution and accuracy of the 3D structures given to DLA-Mutation may influence its performance. Here, we chose to generate 3D models with a high level of precision using the Rosetta backrub protocol, and we explicitly accounted for conformational variability. We showed that averaging over a few tens of conformations improved the discrimination of amino acid classes compared to relying on only one conformation. Future work will more thoroughly investigate the contribution of conformational sampling and the quality of the prediction of mutation-induced binding affinity changes. Alleviating the need for precise models and substantial sampling would improve the scalability of the approach. Another direction for improvement concern the treatment of substitutions to alanine. We found that DLA-Mutation had difficulties in accurately estimating the effects of such mutations and distinguishing them. This trend is not homogeneous across complexes, as illustrated by the good predictions obtained for complexes 3M62, 1CHO and 1JCK (**S 12**). We also showed that the environments around alanine are ambiguous, making it difficult for DLA to recover the amino acid identity. Overall, these results suggest that DLA-Mutation would benefit from a simplified version of the architecture for performing computational alanine scans, which relies only on X-ray crystal structure. Combining DLA-Mutation with alanine scans performed on the wild-type complex would open the way to systematically assess mutational outcomes on protein-protein interactions at a proteome-wide scale.

## Acknowledgements

We thank the Institute for Development and Resources in Intensive Scientific Computing (IDRIS-CNRS) for giving us access to their Jean Zay supercomputer. We thank A. Bonvin for helping us to retrieve the HADDOCK 3D models for the comparison with iSEE.

## Supplementary Information

**Table S 1:** Benchmark datasets of changes of binding affinity upon mutation

| Name | Number of complexes | Number of mutations | Type of point mutations | Source Database |
| --- | --- | --- | --- | --- |
| ZEMu [15] | 65 | 1240 | single+multiple | SKEMPI1 |
| S1102 [20] | 57 | 1102 | single | SKEMPI1 |
| S487 [20] | 56 | 487 | single | SKEMPI2 |
| S645 [54] | 29 | 645 | single | AB-Bind |
| S787 [73] | 24 | 787 | single | AB-Bind |
| S4947 [73] | - | 4947 | single | SKEMPI2 |
| S4169 [59] | 319 | 4169 | single | SKEMPI2 |
| S8338<sup>†</sup> [59] | 319 | 8338 | single | SKEMPI2 |
| S1721 [60] | 147 | 1721 | single+multiple | SKEMPI2 |
| S1402 [75] | 114 | 1402 | single+multiple | SKEMPI1 |
| M1707 [76] | 120 | 1707 | multiple | SKEMPI2 |
| S2003 | 142 | 2003 | single | SKEMPI2 |

<sup>†</sup> S8338 is generated from S4169. It doubles the number of samples by assigning reverse mutation energy changes to the negative values of its original energy values in order to increase the robustness of the predictive method.

### Definition of the interfacial residues

We define interfacial residues as those displaying a change in solvent accessibility between the free (isolated) protein and the complex [39]. We used NACCESS [29] with a probe radius of 1.4Å to compute residue solvent accessibility.

### Building the cubic volumetric map

To build the cubic volumetric map, the atomic coordinates of the input structure are first transformed to a density function [52]. The density *d* at a point 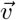 is computed as

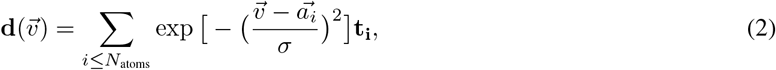

where 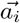 is the position of the *ith* atom, *σ* is the width of the Gaussian kernel set to 1Å, and *t_i_* is a vector of 167 channels corresponding to residue-specific atom types, or 4 channels corresponding to the four amino acid-independent chemical elements (O, C, N and S) (see [52] for a detailed list). The hydrogen atoms are discarded. Then, the density is projected on a 3D grid comprising 24 × 24 × 24 voxels of side 0.8Å. The map is oriented by defining a local frame based on the common chemical scaffold of amino acid residues in proteins [52]. More precisely, for the *nth* residue, the 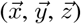 directions and the origin of the cube are defined by the position of the atom N_*n*_, and the directions of *C*_*n*–1_ and *Cα_n_* with respect to N_*n*_. The X-axis is parallel to the vector pointing from C_*n*–1_ to N_*n*_. The Y-axis, perpendicular to the X-axis, is defined in such a way that *Cα_n_* lies in the half-plane O*xy* with *y* > 0. The Z-axis is defined as the vector product *X* × *Y*. The origin of the cube is determined in such a way that N_*n*_ is located at position (6.1Å, 6.6Å, 9.6Å). This choice ensures that all the atoms of the central residue fit in the cube. More details can be found in [52]. This representation is invariant to the global orientation of the structure while preserving information about the atoms and residues relative orientations.

**Table S 2:** Different approaches for the prediction of changes of binding affinity upon mutation.

| Approach | Type | Information | $\Delta\Delta G_{bind}$<br>directly | Train<br>set | Test<br>set | PCC | RMSE<br>($\frac{kcal}{mol}$) |
| --- | --- | --- | --- | --- | --- | --- | --- |
| FLEX [3] | Physics | Structure | - | - | ZEMu | 0.63 | - |
| BindProfX [75] | Physics+Statistics | Structure+Sequence | ✓ | - | S1402 | 0.691 | - |
| iSEE [20] | ML | Structure+Sequence | ✓ | S1102<br>S1102 | -<br>S487 | 0.80*<br>0.25 | 1.41<br>1.32 |
| mCSM-AB [54] | ML | Structure | ✓ | S645 | - | 0.53 | - |
| mCSM-PPI2 [59] | ML | Structure | ✓ | S4169<br>S8338 | -<br>- | 0.76*<br>0.82* | 1.19<br>1.18 |
| mmCSM-PPI [60] | ML | Structure | ✓ | S1721<br>S1721† | -<br>S1721† | 0.87*<br>0.70 | 1.41<br>2.06 |
| TopNetTree | ML | Structure | ✓ | S4947 | - | 0.82* | 1.11 |
|  |  |  |  | S4169 | - | 0.79* | 1.13 |
|  |  |  |  | S8338 | - | 0.85* | 1.11 |
|  |  |  |  | S645 | - | 0.65* | 1.57 |
|  |  |  |  | S4947 | S787 | 0.53 | 1.45 |
| MuPIPR | ML | Sequence | ✓ | S1102 | - | 0.85* | 1.23 |
|  |  |  |  | S1400 | - | 0.88* | 1.32 |
|  |  |  |  | S1102 | S487 | 0.25 | 1.36 |
| GraphPPI | ML | Structure | ✓ | S645 | - | 0.67* | - |
|  |  |  |  | M1707 | - | 0.88* | - |
|  |  |  |  | S645 | - | 0.53§ | - |
|  |  |  |  | M1707 | - | 0.76§ | - |
|  |  |  |  | S645 | - | 0.48¶ | 1.74 |
|  |  |  |  | M1707 | - | 0.73¶ | 2.26 |

\* Mutation-based cross validation, in which a complex (or even the same mutation position of that complex) can be found in different folds. § Leave-one-complex-out cross validation. ¶ Leave-one-structure-out cross validation. † A subset of 1126 mutations used for training/CV and a subset of 595 mutations held out as non-redundant blind test at mutation level. ML: Machine learning, PCC: Pearson Correlation Coefficient, RMSE: Root Mean Squared Error.

### Auxiliary features

For predicting ΔΔ*G_bind_,* we combined the embedding vectors of the volumetric maps with five pre-computed auxiliary features (**Fig. 1C**), among which four describe the wild-type residue:

- a one-hot vector encoding the protein structural region to which it belongs, either the interior (INT), the surface (SUR), or, if it is part of the interface, the support (S or SUP), the core (C or COR), or the rim (R or RIM), as defined in [39]. We directly took the annotations available in the SKEMPI database [31] (see below for a description of the database). We previously demonstrated the usefulness of the S-C-R classification for predicting and analysing protein interfaces with other macromolecules (protein, DNA/RNA) [48, 9, 57, 37].
- its physico-chemical properties (PC, a float value) to be found at interfaces, scaled between 0 and 1 [50].
- its circular variance (CV, a float value) [45, 7] with a sphere radius of 12 Å on the protein structure. For each protein atom, CV measures the density of protein atoms around it within a sphere. The CV of a given residue is obtained by averaging values over its atoms and indicates its degree of burial in the protein. CV values range from 0 to 1 and protruding residues have a value close to 0.
- its conservation level *T_jet_* (a float value) determined by the Joint Evolutionary Trees (JET) method [18]. JET estimates evolutionary conservation by explicitly accounting for the topology of the phylogenetic tree relating the query protein to its homologs.

We used the JET2 package [37] to compute PC, CV and *T_jet_*. We previously showed the usefulness of these properties for detecting protein-protein interfaces and inferring their functions [37]. The fifth feature is specific of the mutation, that is

- a numerical score (a float value) estimating the functional impact of point mutations from multiple sequence alignments computed for single (monomeric) proteins by GEMME [38]. To do this estimation, GEMME combines the conservation levels *T_jet_* with amino acid frequencies and the minimum evolutionary distance between the protein sequence and an homologous protein presenting the mutation.

We built the input multiple sequence alignments for JET and GEMME by performing five iterations of the profile HMM homology search tool Jackhmmer [34, 16] against the UniRef100 database of non-redundant proteins [66] using the EVcouplings framework [27]. We used the default bitscore threshold of 0.5 bit per residue.

### Experimental values for ΔΔ*G_bind_*

We used SKEMPI v2.0 [31], the most complete source for experimentally measured binding affinities of wild-type and mutated protein complexes. It includes the smaller databases AB-Bind, PROXiMATE, and dbMPIKT [21]. In total, it reports measurements for over 7 000 single and multiple point mutations coming from 345 protein complexes, including antibody-antigen (AB/AG) and protease-inhibitor (Pr/PI) assemblies, and assemblies formed between major histocompatibility complex proteins and T-cell receptors (pMHC-TCR). For each entry, corresponding to a single or multiple mutation, the database provides the PDB structure of the wild-type complex, the names of the partners, the binding affinities of the wild-type and mutated complexes, some related experimental measurements, details about the experimental method and conditions, and the structural region of the mutation site(s), either INT, SUR, SUP, COR or RIM [39]. The mutations happening in the interface (SUP, COR, RIM), in particular in the core (COR), induce bigger changes in binding affinity than the ones located in the non-interacting surface (SUR) or the interior (INT) of the protein (**Fig. S 2)**. Overall, we observed a tendency for the mutations to be deleterious rather than beneficial. The most impactful single-point mutation is located in the complex 1CHO with ΔΔ*G_bind_* = 8.802 kcal/mol. This rich body of annotations helps us to analyze our results and identify the weak and strong points of DLA-mutation by evaluating its performance with respect to different classifications.

We restricted our experiments to the entries for which the binding affinity of the wild-type and mutant complexes were determined using a reliable experimental method, namely ITC, SPR, FL, or SP, as done in [70]. This first filtering step led to 4 974 entries associated with 255 protein complexes. We retained 4 634 entries from 245 complexes by excluding mutation entries with ambiguous free energy or without energy change. We then focused only on 3 393 single-mutation entries coming from 222 complexes. After removing duplicated entries (a protein complex with the same mutations), we remained with 2 975 mutations. We finally randomly selected a subset of 2 003 mutations associated with 142 complexes. We call this subset *S2003.*

### Protein-protein complex 3D structures

We created two databases of protein-protein complex 3D structures, namely *PDBInter* and *S2003-3D,* for training and validation purposes. PDBInter was curated from the Protein Data Bank (PDB) [5] and thus contains only experimental structures. S2003-3D was generated using the “backrub” protocol implemented in Rosetta [64] and thus contains only 3D models.

#### PDBInter

We downloaded all PDB biological assemblies (June 2020 release) from the FTP archive rsync.wwpdb.org::ftp/data/biounitrsync.wwpdb.org::ftp/data/biounit. We discarded the entries with more than 100 chains or with a resolution lower than 5Å. We also removed the protein chains smaller than 20 residues or with more than 20% of unknown residues. We then redundancy-reduced the resulting dataset using annotations from the SCOPe database [19, 8]. The 5 055 protein complex structures that were finally retained do not share any family level similarity between them according to the SCOPe hierarchy.

#### S2003-3D

We generated conformational ensembles for the wild-type and mutated complexes from S2003. We followed a modeling protocol similar to that reported in [3]. It relies on the backrub method [64] for sampling side chain and backbone conformational changes. Our goal was to accurately mimic and explore the fluctuations around a native state. The protocol unfolds in two optimization steps carried out on the side chains and the backbone (**Fig. S 1)**:

1. for the backbone and the side chains, it applies quasi-Newton minimization for continuous optimization of torsion angles: Φ, Ψ, χ_1_, χ_2_, χ_3_, etc.
2. for the side chains only, it performs Monte Carlo simulation with the backbone-based side-chain rotamer library of Dunbrack [61] for discrete combinatorial rotamer optimization, also known as repacking.

**Figure S 1:**
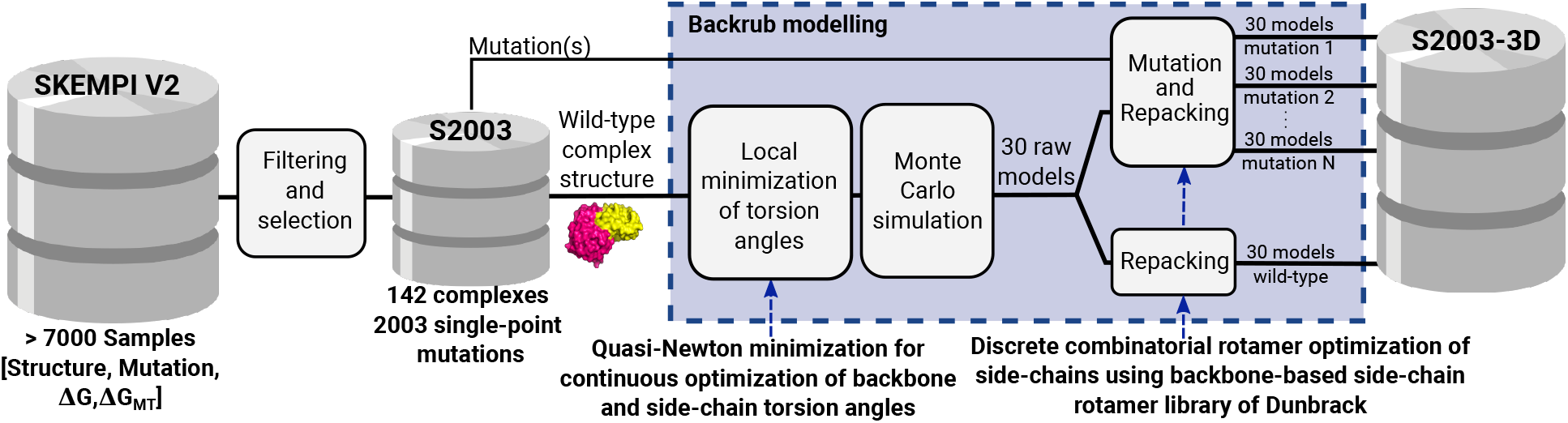
Pipeline for the generation of mutated complexes with backrub. After filtering the SKEMPI v2.0 database, we retained 2003 single point-mutations for 142 complexes (S2003). A wild-type structure undergoes a local minimisation of backbone and side-chain torsion angles followed by a Monte Carlo simulation step. We applied it to produce 30 models for each mutated structure and 30 for the wild-type. This process is followed by a repacking step applied to wild-type and mutation models. For the mutation positions at the interface of each model, we compute the associated cubic volumetric maps.

### Evaluation of ssDLA on the validation set from PDBInter

To visualise the performance of the model, we generated logos from pseudo alignments of 20 columns corresponding to the 20 amino acids. In the column corresponding to the amino acid *a_i_*, the frequency of occurrence of each amino acid *a_j_* corresponds to the propensity of ssDLA to predict *a_j_* when the true central residue of the input cube is *a_i_*. Note that the propensity is computed by counting the number of times *aj* has maximum probability score among the 20 candidate amino acids. If some amino acid was never predicted, we simply put a gap character.

### Different experimental setups for the supervised prediction of ΔΔ*G_bind_*

We experimented different setups of supervised learning by using different combinations of auxiliary features and different initialisation schemes for the network weights. In the basic set up, the only auxiliary feature we used was the structural region of the wild-type residue (SR). We previously showed that this information significantly contributes to the performance of the DLA framework [48]. On top of that, we also considered evolutionary information, by using the *GEMME* scores (*SR-GEMME*) or the *T_jet_* conservation levels (*SR-Tjet*). In its most complete form, DLA-mutation combines SR, *GEMME* scores, *T_jet_*, and descriptors of the buriedness (CV) and physico-chemical properties (PC) of the wild-type residue (All). For the network weights, we either started from the weights of the pre-trained ssDLA (*fine tuning*) or randomly initialised them. For each mutation, DLA-Mutation considers 30 pairs of cubes extracted from the mutation site of the associated 30 backrub models. The predicted ΔΔ*G_bind_* is an average over all 30 models.

**Table S 3:** Seven classes of amino acids.

| Class name | Description | Amino acid(s) | Representative color |
| --- | --- | --- | --- |
| ARO | Aromatic | F, W, Y, H | Green |
| CAST | Hydroxyl-containing and Alanine | C, A, S, T | Black |
| PHOB | Aliphatic hydrophobic | I, L, M, V | Red |
| POS | Positively charged | K, R | Purple |
| POL-N | Polar and negatively charged | N, Q, D, E | Blue |
| GLY | Glycine | G | Gray |
| PRO | Proline | P | Orange |

### Training and evaluation of downstream tasks: prediction of residue- and interface-level properties

We generated embedding vectors (*e_k_*) for all the cubes representing a given input interface. For training purposes, we redundancy-reduced the set of 142 complexes from *S2003* based on a 30% sequence identity cutoff. We then performed a 50/50 split at the cluster level. This resulted into 85 train and 57 test complexes for the first, residue-based, task. As training and testing samples, we considered:

- either all interfacial residues (4710 residues for train and 3397 residues for test) extracted from the X-ray crystal structures of *S2003;*
- or only the residues belonging to the positions with mutation from S2003 (1700 residues for train and 303 residues for test) extracted from the wild-type backrub models of *S2003-3D.* We performed two experiments here: (i) pick up one backrub model at random (out of 30) to generate the input cubes, (ii) average the embedding vectors computed for a given interfacial residue over the 30 backrub models.

For the second, interface-based task, due to missing annotations, we used only 22 train and 52 test complexes. We computed the average embedding vector over all interfacial residues before giving it to the classifier. In addition, we focused only on the X-ray crystal structures. In both tasks, the number of epochs depended on the size of train set and the learning rate (0.00001). We stopped the training when the validation loss converged to a steady value (**Fig. S 8)**.

**Table S 4:** Weights of amino acids classes in self-supervised learning.

| Amino acid | Weights |  |
| --- | --- | --- |
|  | 167 channels | 4 channels |
| A | 0.768 | 0.768 |
| C | 4.100 | 4.100 |
| D | 0.901 | 0.751 |
| E | 0.724 | 0.724 |
| F | 1.117 | 1.117 |
| G | 0.825 | 0.825 |
| H | 1.747 | 1.747 |
| I | 0.920 | 0.920 |
| K | 0.904 | 0.904 |
| L | 0.529 | 0.529 |
| M | 2.088 | 2.088 |
| N | 1.170 | 1.170 |
| P | 0.856 | 0.856 |
| Q | 1.182 | 1.182 |
| R | 0.717 | 0.717 |
| S | 0.885 | 0.885 |
| T | 0.920 | 0.920 |
| V | 0.817 | 0.817 |
| W | 2.897 | 2.897 |
| Y | 1.109 | 1.109 |

**Figure S 2:**
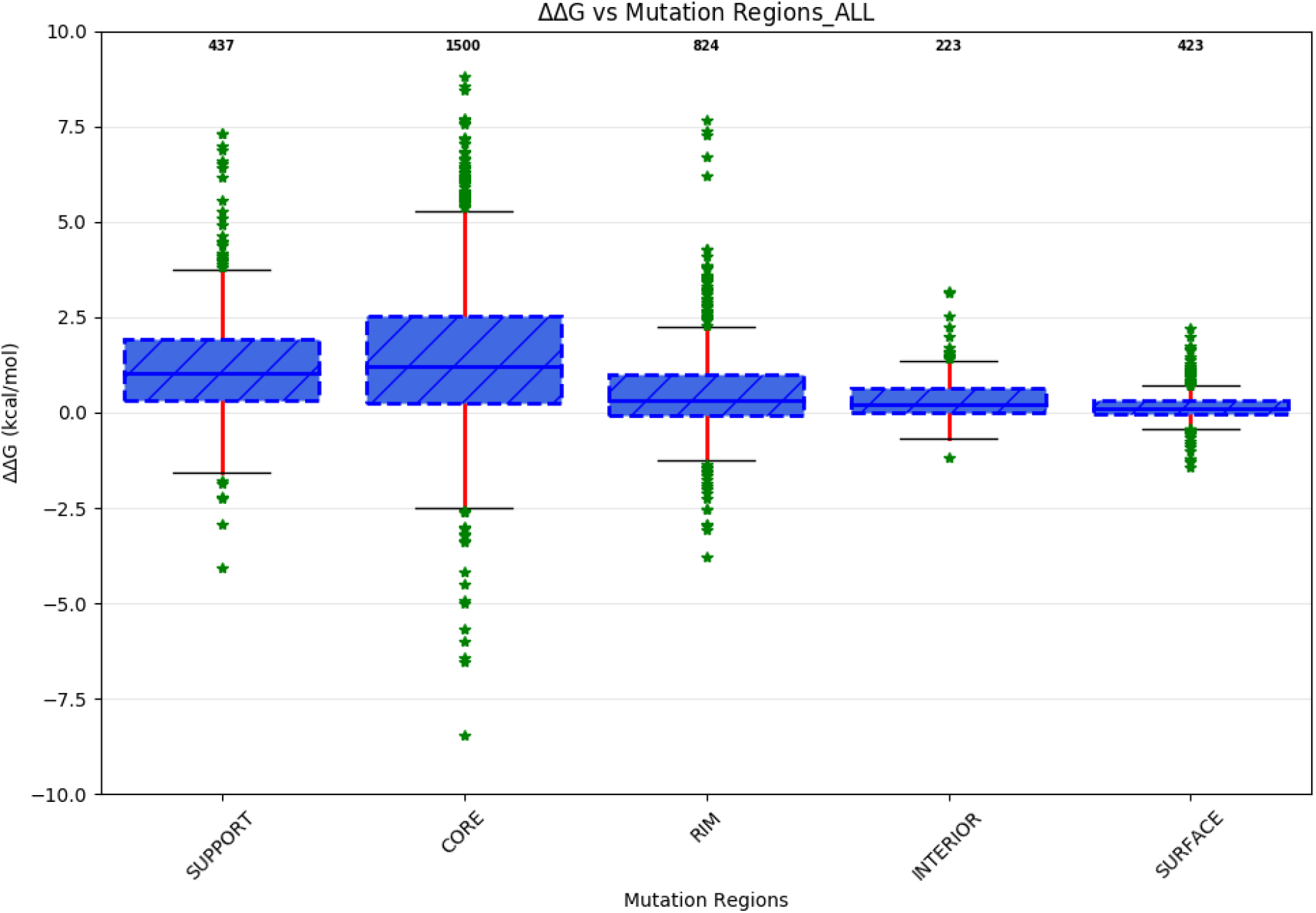
Distribution of ΔΔ*G_bind_* values in SKEMPI v2. We focus here only on the single point mutations. The different distributions correspond to the different protein structural regions: COR (1500 mutations), SUP (437 mutations), RIM (824 mutations), INT (223 mutations), and SUR (423 mutations).

**Figure S 3:**
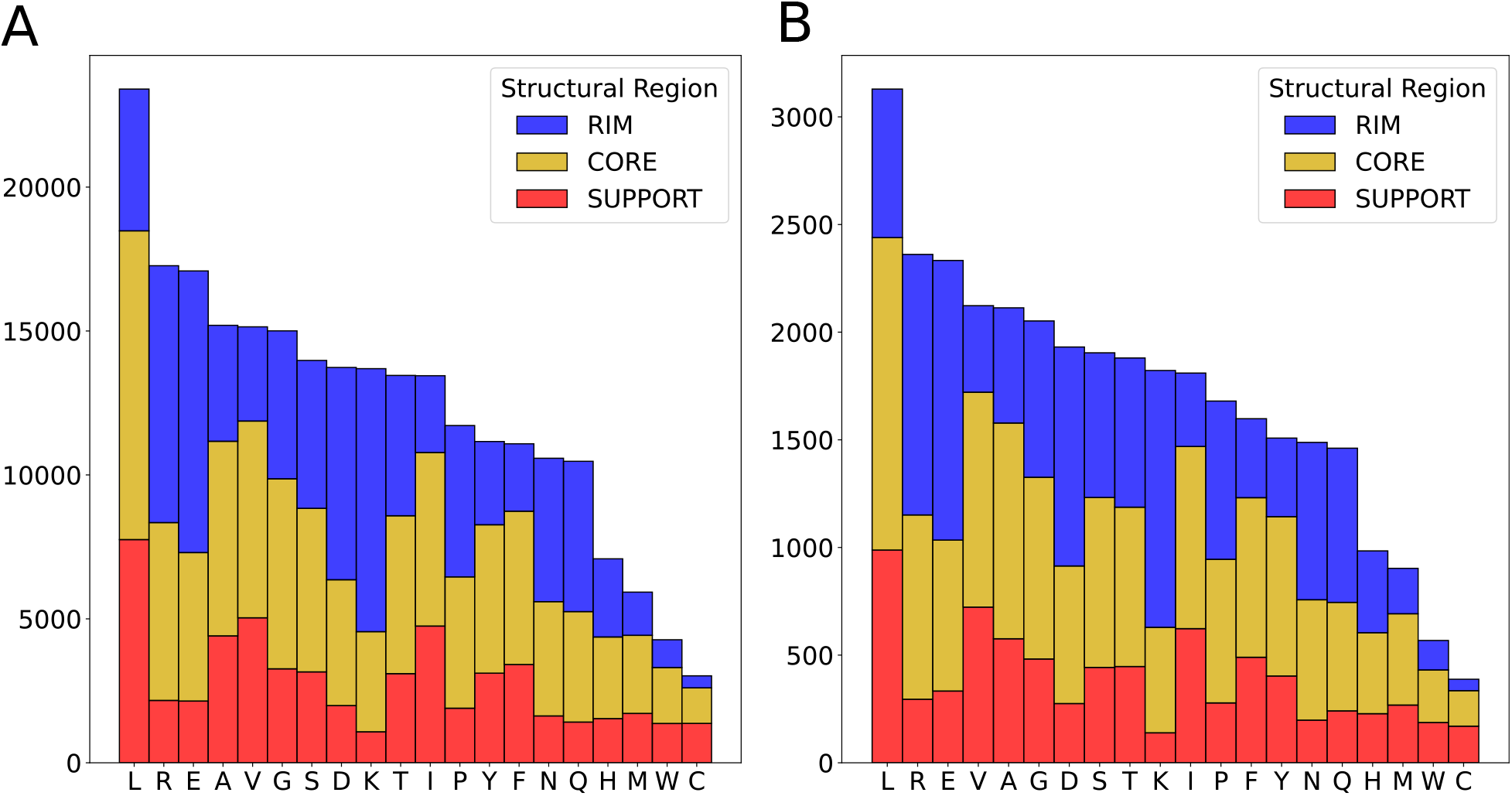
Frequency of interfacial amino acids in PDBInter. For each amino acid type, we report the number of times it appears in the core, the rim, or the support of the interface. **A** train set. **B.** Validation set.

**Figure S 4:**
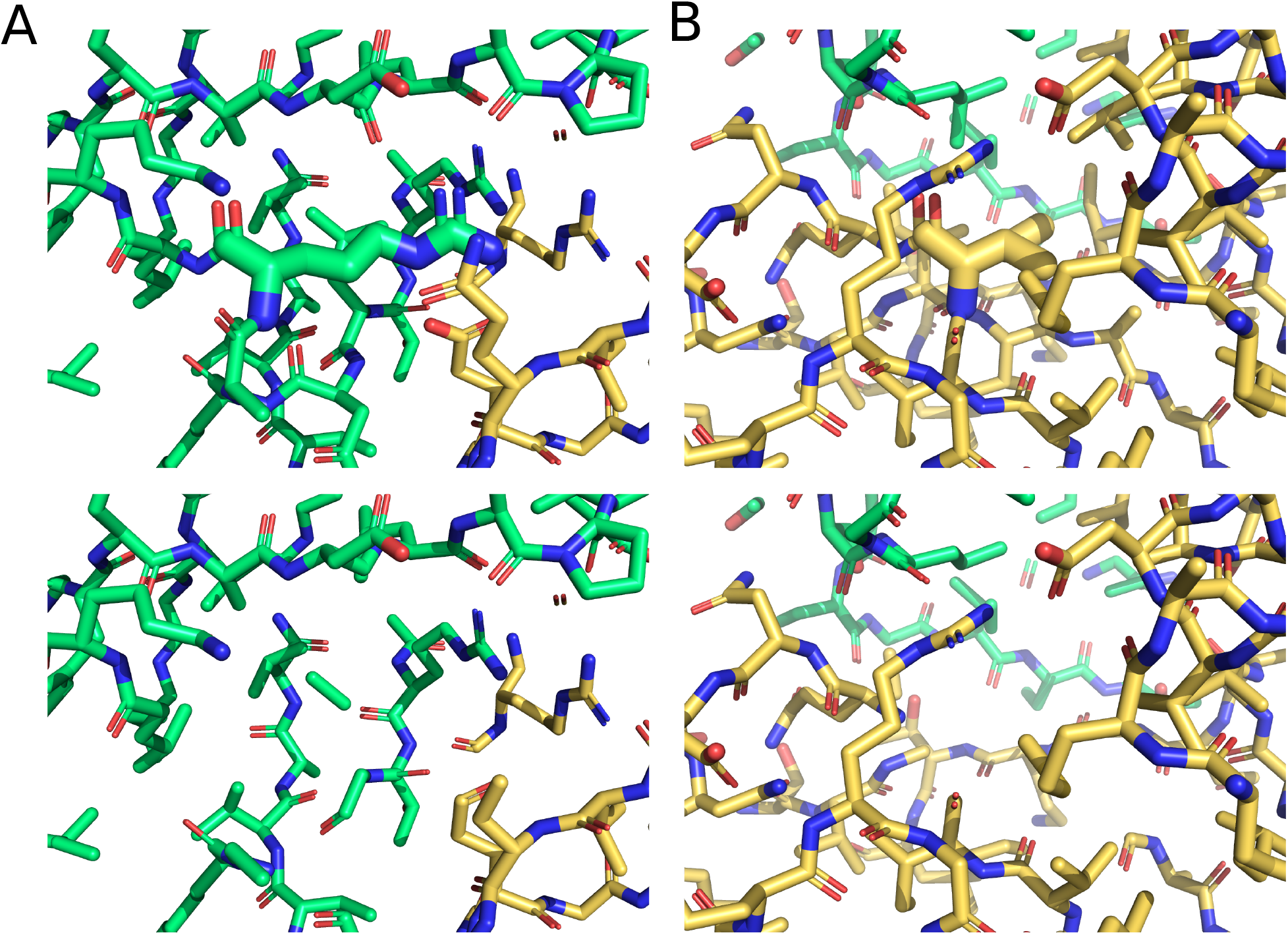
An example of masking with a sphere of 5 Å. Masking atoms inside a spherical volume of radius 5 Å randomly centered on the central interfacial residues (**A**) arginine or (**B**) isoleucine. Top is the intact local environment and bottom is the masked one. Both interfacial residues belong to the same interface.

**Figure S 5:**
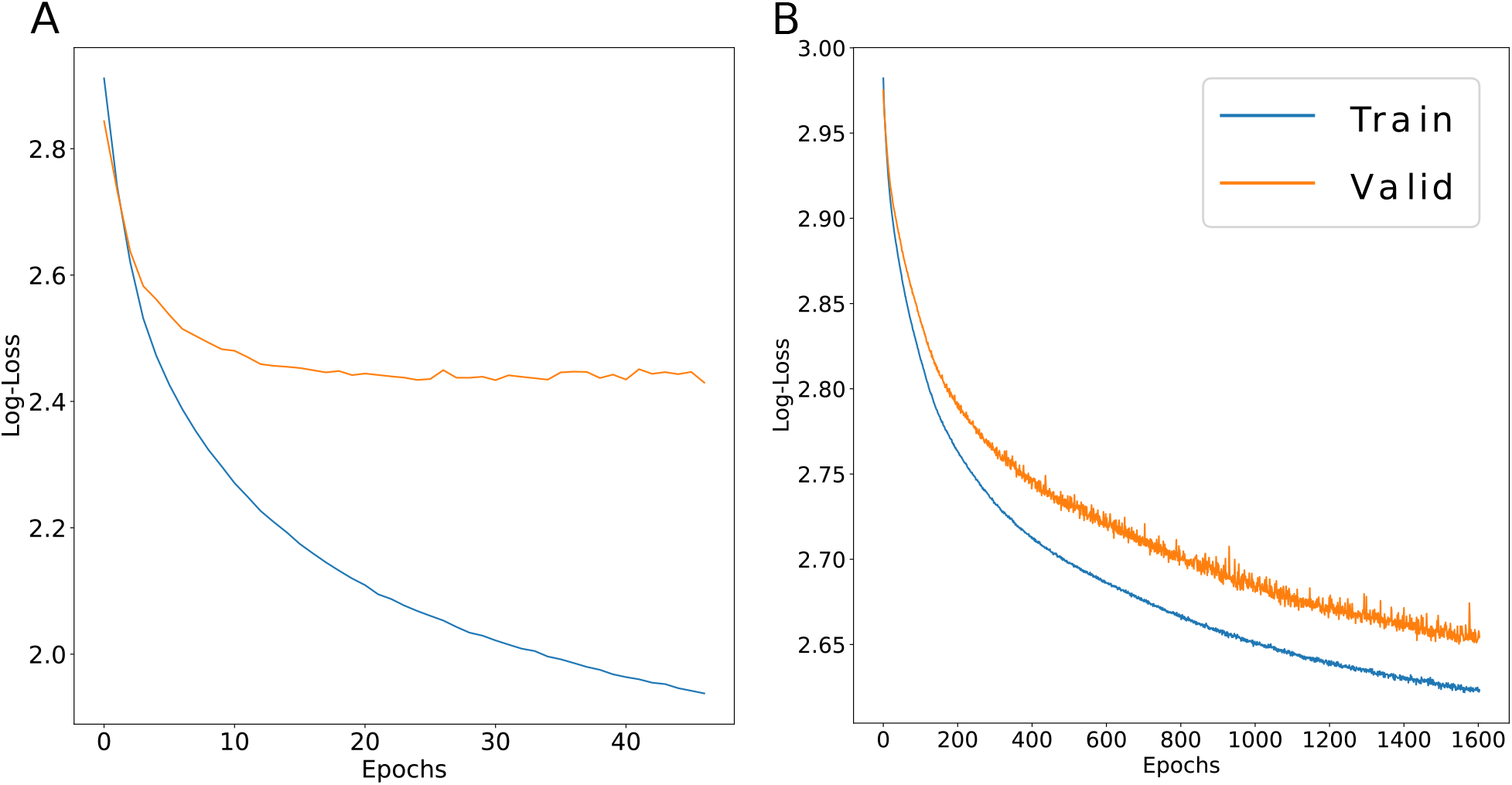
Train and validation loss curves of ssDLA. The x-axis is the number of epochs and the y-axis is the log-loss (categorical cross-entropy). (**A**) Default ssDLA model, where we used 167 channels corresponding to 167 amino acid-specific atom types (see [52] for a detailed list). (**B**) Simplified model where we considered only 4 atom types (C, N, O, S).

**Figure S 6:**
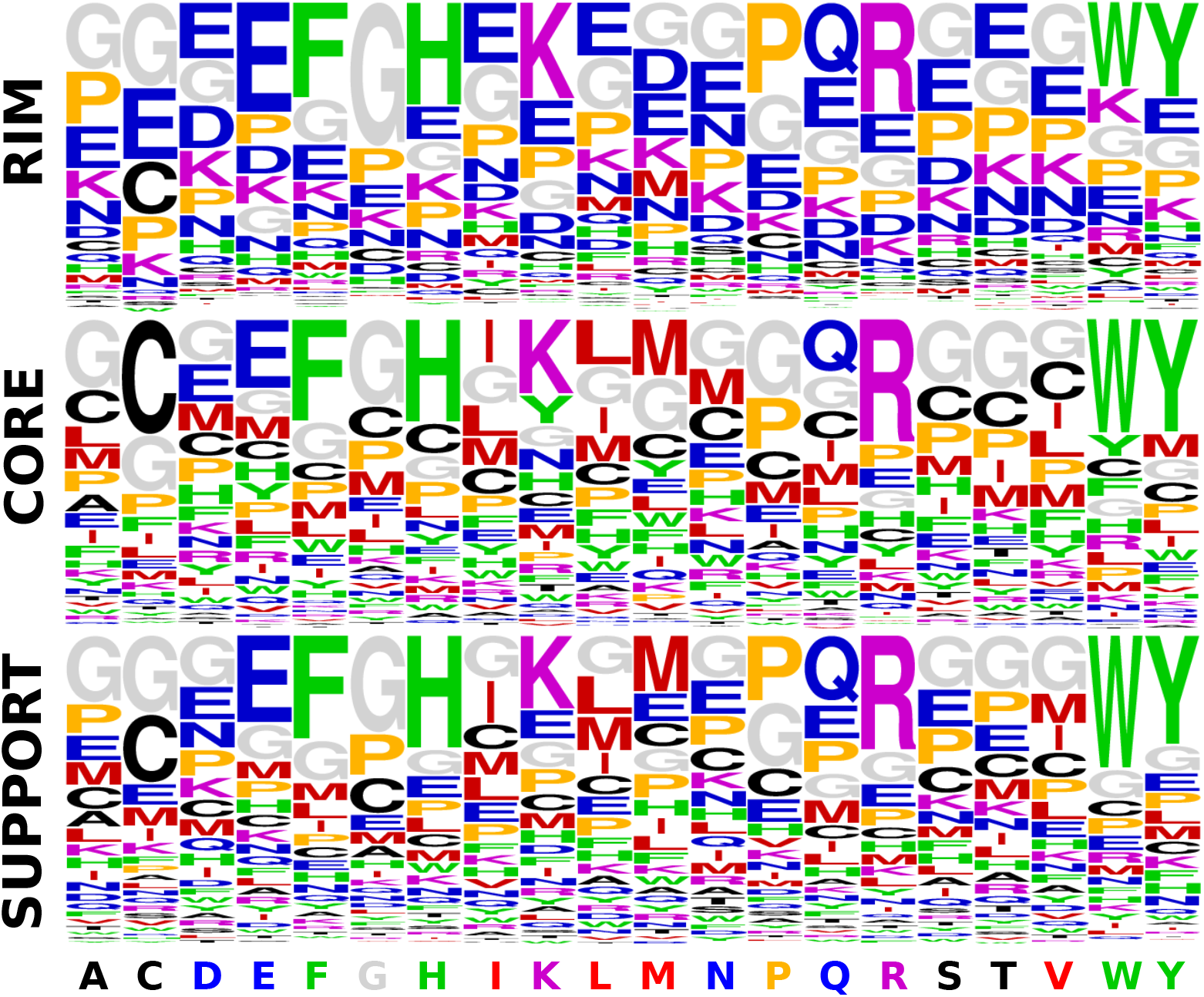
Performance of simplified ssDLA model with 4 channels. The predictive power of simplified ssDLA model where we considered only 4 atom types (C, N, O, S) is evaluated on the validation set of *PDBInter*. The three logos represent the propensities of each amino acid to be predicted (having maximum score in the output layer), depending on the true amino acid (x-axis) and on its structural region (see *Methods*). Amino acids are colored based on seven similarity classes: ARO (F, W, Y, H) in green, CAST (C, A, S, T) in black, PHOB (I, L, M, V) in red, POS (K, R) in purple, POL-N (N, Q, D, E) in blue, GLY (G) in gray and PRO (P) in orange (see *Methods*).

**Figure S 7:**
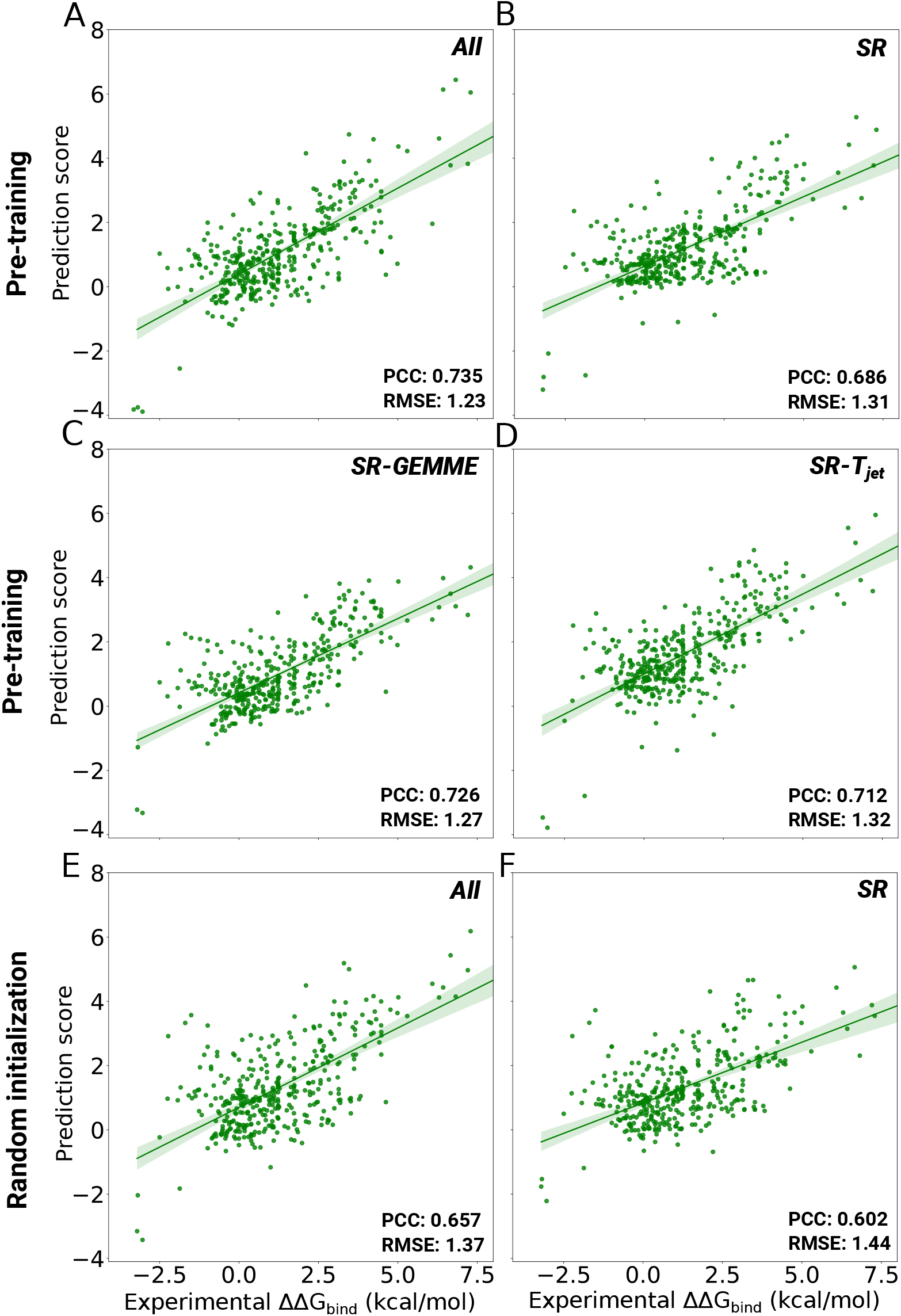
The predictive performance of six experimental setups on a test set of 391 mutations from 32 unseen protein complexes (randomly selected from S2003 dataset) with a complex-based train and test split. **A-D.** The training process fine-tunes the weights of the pre-trained model ssDLA and includes *All* (**A**), *SR* (**B**), *SR-GEMME* (**C**) or *SR-Tjet* (**D**) features. **E-F.** Training starts from randomly initialized weights with *All* (**E**) or *SR* (**F**) features.

**Figure S 8:**
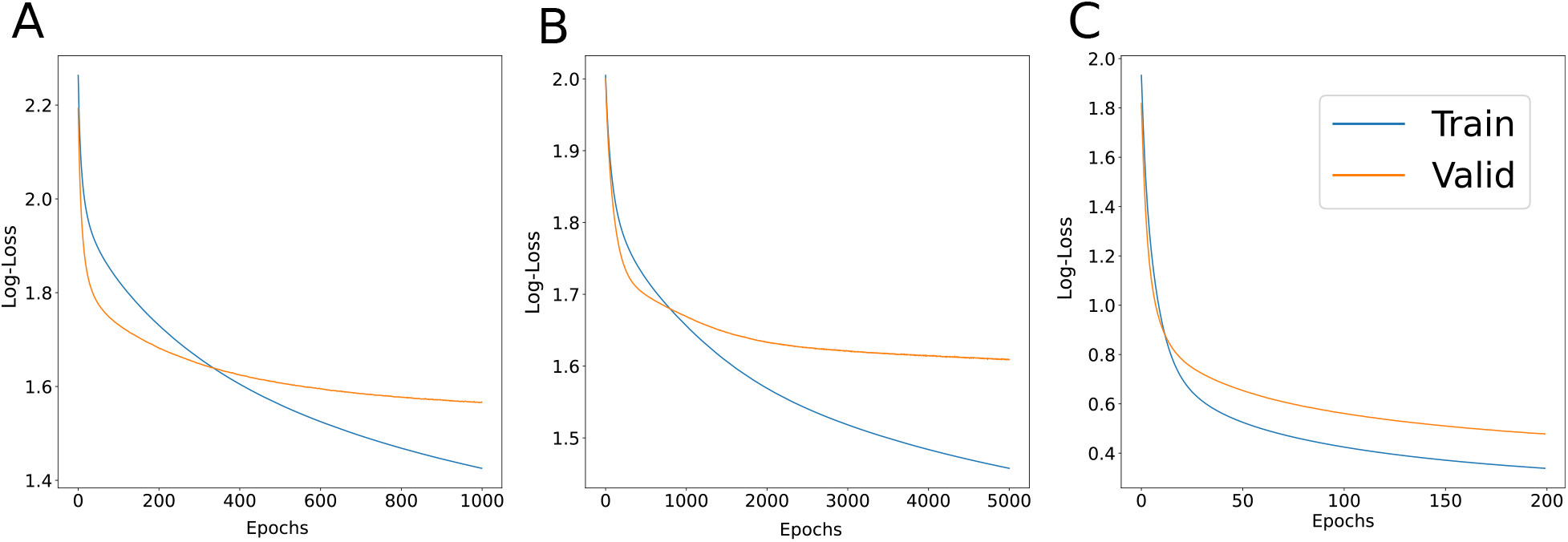
Train and validation loss curves of the downstream task: prediction of amino acid classes. x-axis is the epochs and the y-axis is the loss function (categorical cross-entropy). **A-B.** Embedding vectors extracted by default ssDLA model (167 channels, **A**) or by simplified model (4 channels, **B**) from X-ray crystal structures of S2003. **C.** Embedding vectors extracted by default ssDLA from wild-type backrub models of S2003.

**Figure S 9:**
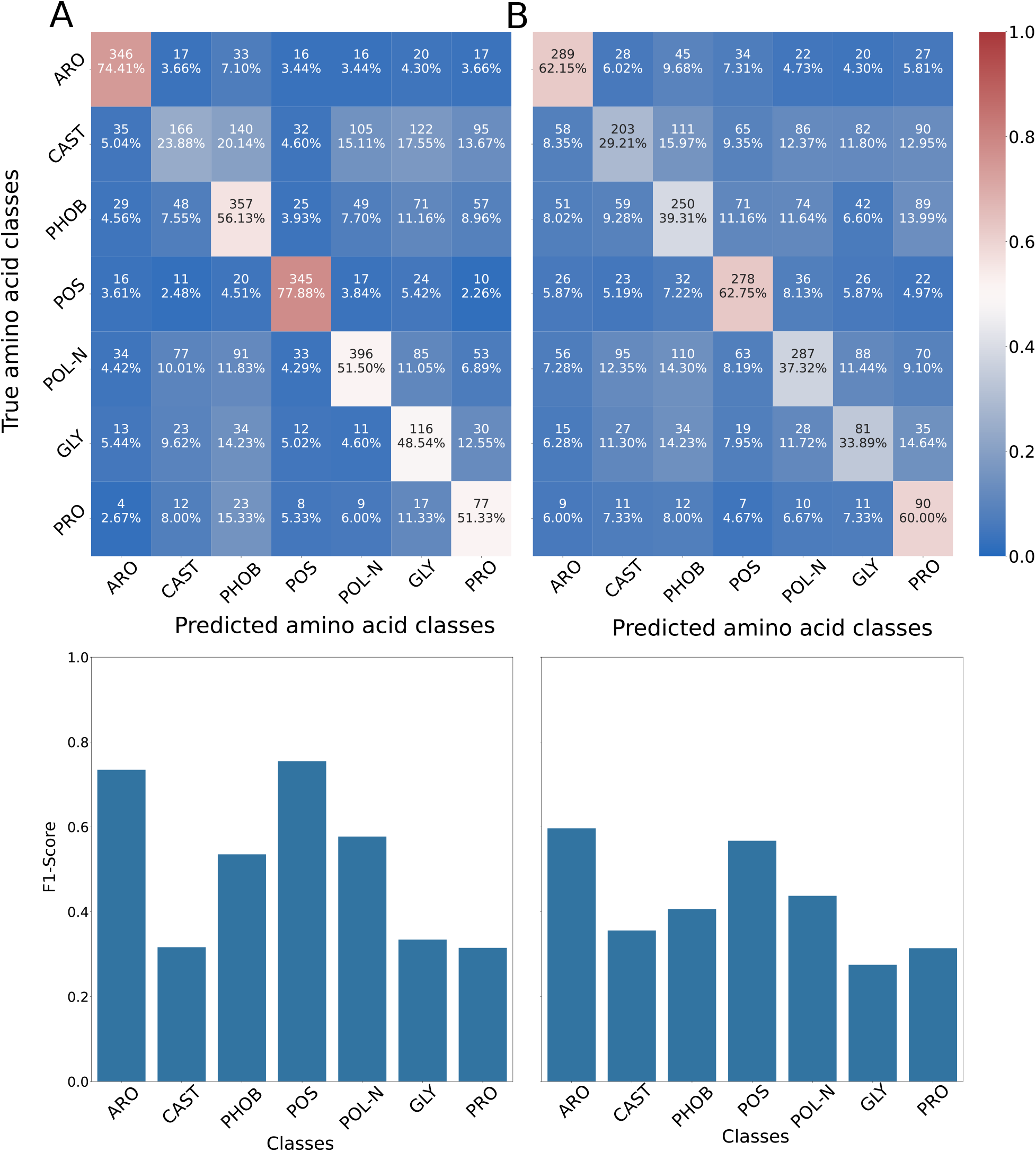
Prediction of the amino acid classes for the interfacial residues using embedding vectors extracted by ssDLA. Train and test were performed on all the interfacial residues of the X-ray crystal structures of S2003. The confusion matrix and per-class F1-scores for the embedding vectors extracted by default ssDLA (167 channels, **A**) or simplified version (4 channels, **B**). In the confusion matrices the percentage values and the colors indicate recall.

**Figure S 10:**
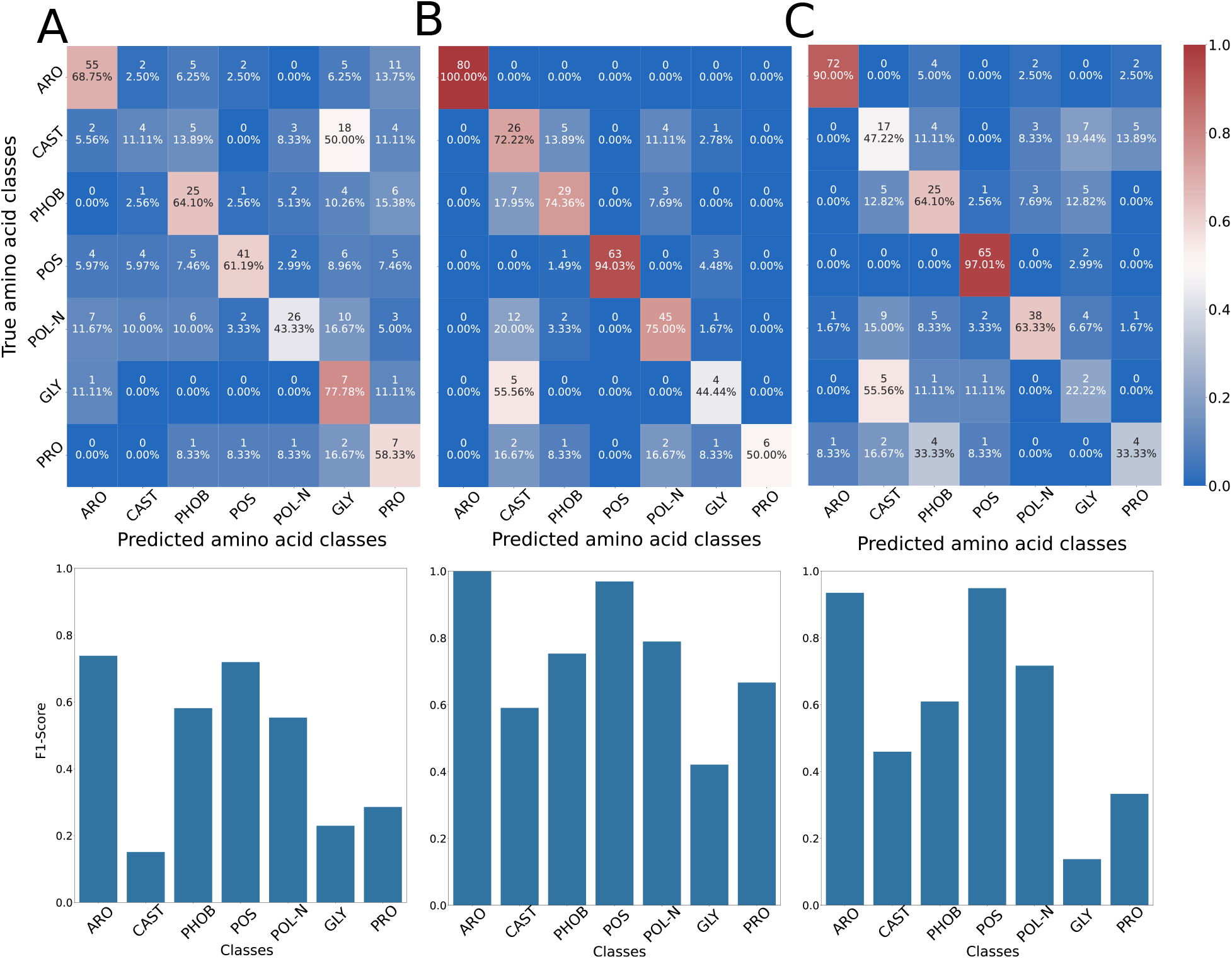
Prediction of the amino acid classes for the mutation sites using embedding vectors extracted by ssDLA. Train and test were performed on the subset of mutation sites of the X-ray crystal and wild-type backrub structures of S2003. **A.** The confusion matrix and per-class F1-scores for the embedding vectors of X-ray structures extracted by default ssDLA (167 channels). **B-C.** The confusion matrix and per-class F1-scores for the embedding vectors of wild-type backrubs extracted by default ssDLA with two aggregating schemes: averaging over backrub models (**B**) or choosing a single backrub model (**C**). In the confusion matrices the percentage values and the colors indicate recall.

**Figure S 11:**
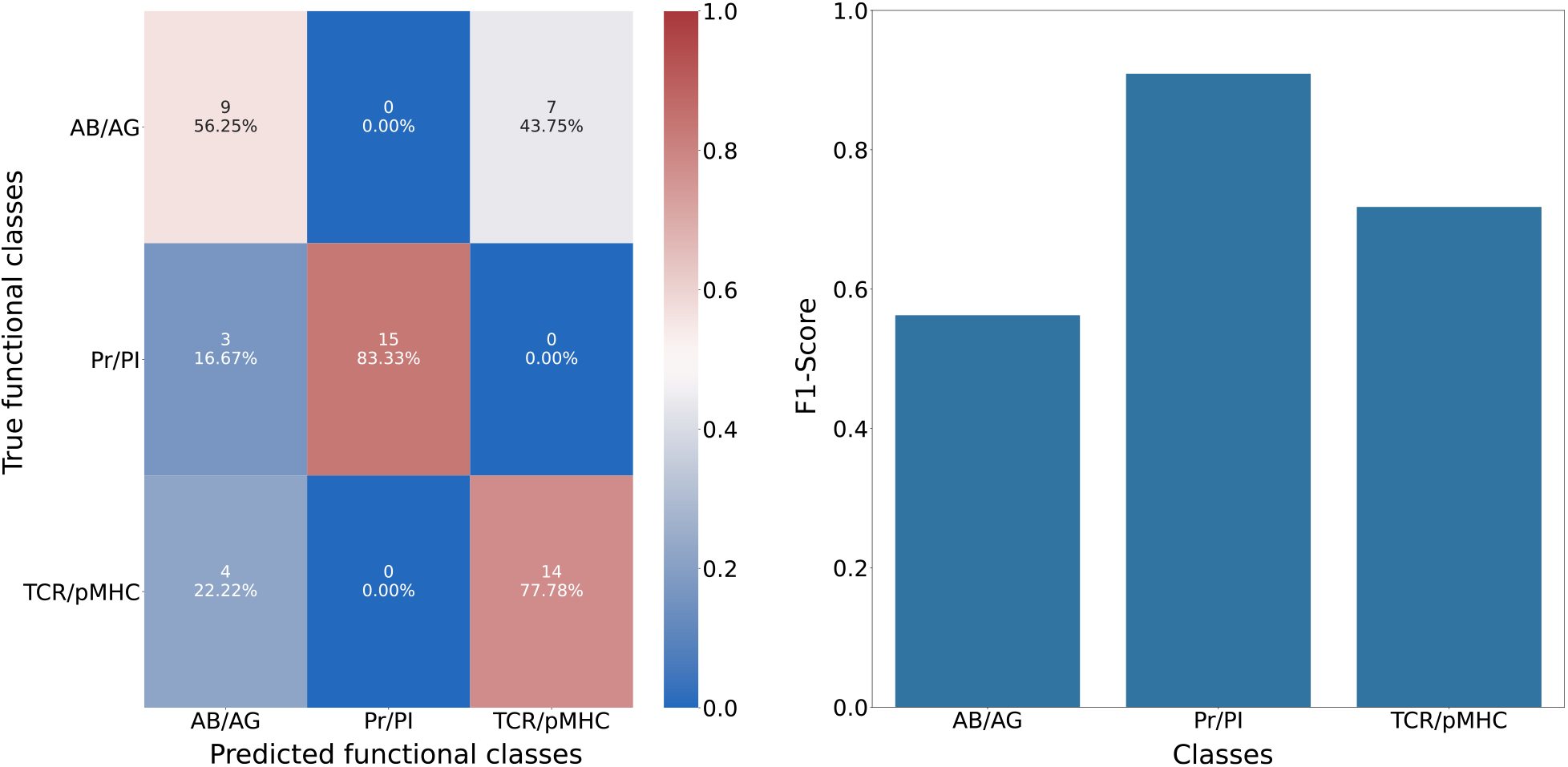
Prediction of the interaction functional classes using embedding vectors extracted by ssDLA. Train and test were performed on the X-ray crystal structures of S2003. **A-B.** The confusion matrix (**A**) and per-class F1-scores (**B**) for the embedding vectors extracted by default ssDLA (167 channels). In the confusion matrices the percentage values and the colors indicate recall. The aggregating scheme for each complex is the averaging of embedding vectors over is interfacial residues.

**Figure S 12:**
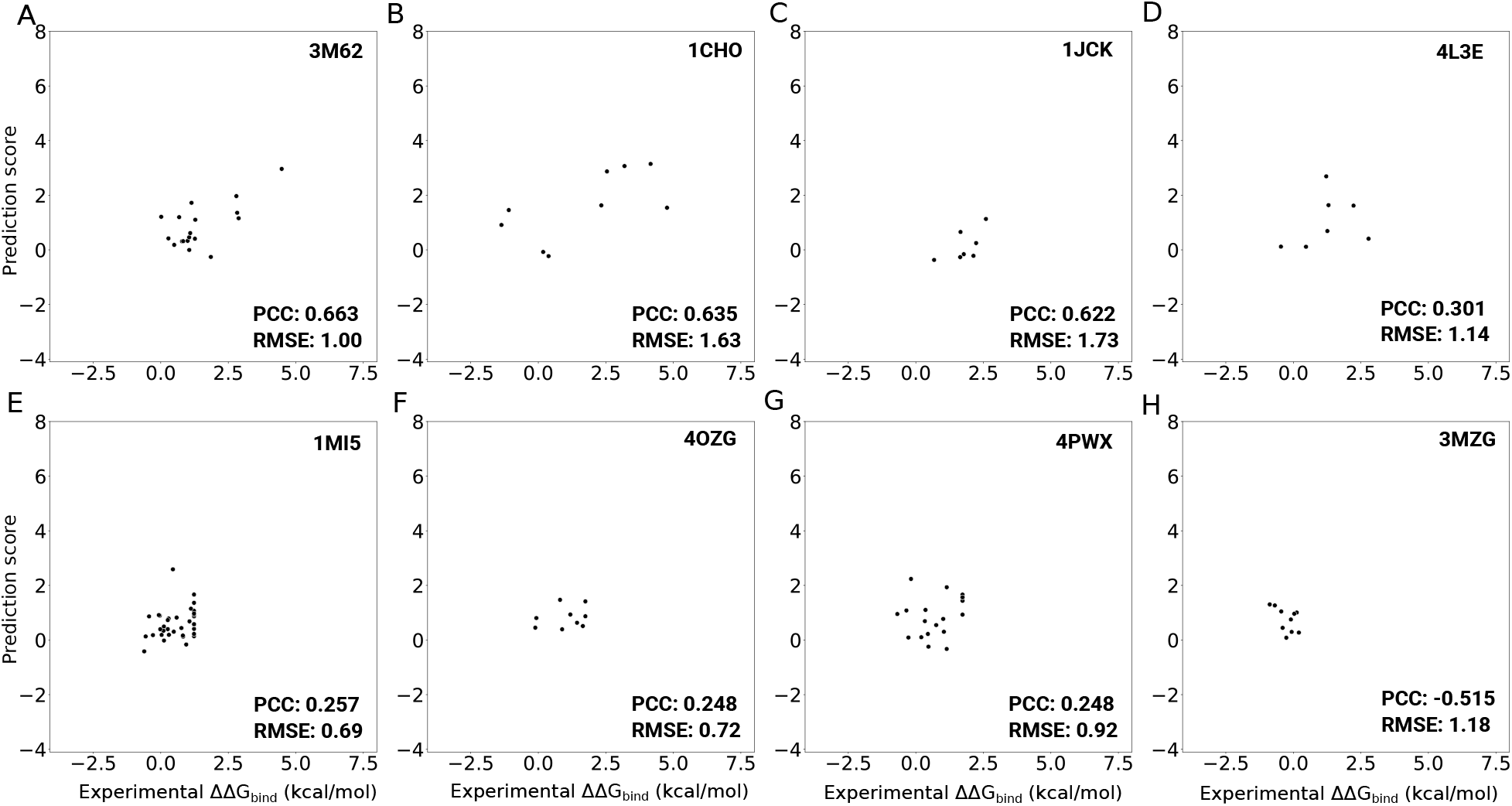
DLA-Mutation predictions for mutations to Alanine separated by different complexes. Predictions are obtained with the DLA-Mutation architecture with a fine-tuned pre-trained model and *All* auxiliary features. **A.** 3M62, **B.** 1CHO, **C.** 1JCK, **D.** 4L3E, **E.** 1MI5, **F.** 4OZG, **G.** 4PWX, **H.** 3MZG.

## References

[1] Martin Abadi, Paul Barham, Jianmin Chen, Zhifeng Chen, Andy Davis, Jeffrey Dean, Matthieu Devin, Sanjay Ghemawat, Geoffrey Irving, Michael Isard, Manjunath Kudlur, Josh Levenberg, Rajat Monga, Sherry Moore, Derek G. Murray, Benoit Steiner, Paul Tucker, Vijay Vasudevan, Pete Warden, Martin Wicke, Yuan Yu, and Xiaoqiang Zheng. TensorFlow: A System for Large-Scale Machine Learning. pp. 265–283, 2016.

[2] Namrata Anand, Raphael Eguchi, Irimpan I. Mathews, Carla P. Perez, Alexander Derry, Russ B. Altman, and Po-Ssu Huang. Protein sequence design with a learned potential. Nature Communications, 13(1):746, February 2022.

[3] Kyle A. Barlow, Shane Ó Conchúir, Samuel Thompson, Pooja Suresh, James E. Lucas, Markus Heinonen, and Tanja Kortemme. Flex ddG: Rosetta Ensemble-Based Estimation of Changes in Protein–Protein Binding Affinity upon Mutation. The Journal of Physical Chemistry B, 122(21):5389–5399, May 2018.

[4] Tristan Bepler and Bonnie Berger. Learning the protein language: Evolution, structure, and function. Cell Systems, 12(6):654–669.e3, June 2021.

[5] H. M. Berman, T. Battistuz, T. N. Bhat, W. F. Bluhm, P. E. Bourne, K. Burkhardt, Z. Feng, G. L. Gilliland, L. Iype, S. Jain, P. Fagan, J. Marvin, D. Padilla, V. Ravichandran, B. Schneider, N. Thanki, H. Weissig, J. Westbrook, and C. Zardecki. The Protein Data Bank. Acta Crystallographica Section D: Biological Crystallography, 58(6):899–907, June 2002.

[6] Lasse M. Blaabjerg, Maher M. Kassem, Lydia L. Good, Nicolas Jonsson, Matteo Cagiada, Kristoffer E. Johansson, Wouter Boomsma, Amelie Stein, and Kresten Lindorff-Larsen. Rapid protein stability prediction using deep learning representations, August 2022.

[7] Nicoletta Ceres, Marco Pasi, and Richard Lavery. A Protein Solvation Model Based on Residue Burial. Journal of Chemical Theory and Computation, 8(6):2141–2144, June 2012.

[8] John-Marc Chandonia, Lindsey Guan, Shiangyi Lin, Changhua Yu, Naomi K Fox, and Steven E Brenner. SCOPe: improvements to the structural classification of proteins – extended database to facilitate variant interpretation and machine learning. Nucleic Acids Research, 50(D1):D553–D559, January 2022.

[9] Flavia Corsi, Richard Lavery, Elodie Laine, and Alessandra Carbone. Multiple protein-DNA interfaces unravelled by evolutionary information, physico-chemical and geometrical properties. PLOS Computational Biology, 16(2):e1007624, February 2020.

[10] Pau Creixell, Erwin M. Schoof, Craig D. Simpson, James Longden, Chad J. Miller, Hua Jane Lou, Lara Perryman, Thomas R. Cox, Nevena Zivanovic, Antonio Palmeri, Agata Wesolowska-Andersen, Manuela Helmer-Citterich, Jesper Ferkinghoff-Borg, Hiroaki Itamochi, Bernd Bodenmiller, Janine T. Erler, Benjamin E. Turk, and Rune Linding. Kinome-wide Decoding of Network-Attacking Mutations Rewiring Cancer Signaling. Cell, 163(1):202–217, September 2015.

[11] J. Dauparas, I. Anishchenko, N. Bennett, H. Bai, R. J. Ragotte, L. F. Milles, B. I. M. Wicky, A. Courbet, R. J. de Haas, N. Bethel, P. J. Y. Leung, T. F. Huddy, S. Pellock, D. Tischer, F. Chan, B. Koepnick, H. Nguyen, A. Kang, B. Sankaran, A. K. Bera, N. P. King, and D. Baker. Robust deep learning based protein sequence design using ProteinMPNN, June 2022.

[12] Alessia David, Rozami Razali, Mark N. Wass, and Michael J.E. Sternberg. Protein–protein interaction sites are hot spots for disease-associated nonsynonymous SNPs. Human Mutation, 33(2):359–363, 2012.

[13] Alessia David and Michael J. E. Sternberg. The Contribution of Missense Mutations in Core and Rim Residues of Protein–Protein Interfaces to Human Disease. Journal of Molecular Biology, 427(17):2886–2898, August 2015.

[14] Jacob Devlin, Ming-Wei Chang, Kenton Lee, and Kristina Toutanova. BERT: Pre-training of Deep Bidirectional Transformers for Language Understanding. In Proceedings of the 2019 Conference of the North American Chapter of the Association for Computational Linguistics: Human Language Technologies, Volume 1 (Long and Short Papers), pp. 4171–4186, Minneapolis, Minnesota, June 2019.

[15] Daniel F. A. R. Dourado and Samuel Coulbourn Flores. Modeling and fitting protein-protein complexes to predict change of binding energy. Scientific Reports, 6:25406, May 2016.

[16] Sean R. Eddy. Accelerated Profile HMM Searches. PLOS Computational Biology, 7(10):e1002195, October 2011.

[17] Ahmed Elnaggar, Michael Heinzinger, Christian Dallago, Ghalia Rehawi, Yu Wang, Llion Jones, Tom Gibbs, Tamas Feher, Christoph Angerer, Martin Steinegger, Debsindhu Bhowmik, and Burkhard Rost. ProtTrans: Towards Cracking the Language of Lifes Code Through Self-Supervised Deep Learning and High Performance Computing. IEEE Transactions on Pattern Analysis and Machine Intelligence, pp. 1–1, 2021.

[18] Stefan Engelen, Ladislas A. Trojan, Sophie Sacquin-Mora, Richard Lavery, and Alessandra Carbone. Joint Evolutionary Trees: A Large-Scale Method To Predict Protein Interfaces Based on Sequence Sampling. PLOS Computational Biology, 5(1):e1000267, January 2009.

[19] Naomi K. Fox, Steven E. Brenner, and John-Marc Chandonia. SCOPe: Structural Classification of Proteins—extended, integrating SCOP and ASTRAL data and classification of new structures. Nucleic Acids Research, 42(D1):D304–D309, January 2014.

[20] Cunliang Geng, Anna Vangone, Gert E. Folkers, Li C. Xue, and Alexandre M. J. J. Bonvin. iSEE: Interface structure, evolution, and energy-based machine learning predictor of binding affinity changes upon mutations. Proteins: Structure, Function, and Bioinformatics, 87(2):110–119, 2019.

[21] Cunliang Geng, Li C. Xue, Jorge Roel-Touris, and Alexandre M. J. J. Bonvin. Finding the ddG spot: Are predictors of binding affinity changes upon mutations in protein–protein interactions ready for it? WIREs Computational Molecular Science, 9(5):e1410, 2019.

[22] Mileidy W. Gonzalez and Maricel G. Kann. Chapter 4: Protein Interactions and Disease. PLOS Computational Biology, 8(12):e1002819, December 2012.

[23] Raphael Guerois, Jens Erik Nielsen, and Luis Serrano. Predicting Changes in the Stability of Proteins and Protein Complexes: A Study of More Than 1000 Mutations. Journal of Molecular Biology, 320(2):369–387, July 2002.

[24] Ge-Fei Hao, Guang-Fu Yang, and Chang-Guo Zhan. Structure-based methods for predicting target mutation-induced drug resistance and rational drug design to overcome the problem. Drug Discovery Today, 17(19):1121–1126, October 2012.

[25] Yehouda Harpaz, Mark Gerstein, and Cyrus Chothia. Volume changes on protein folding. Structure, 2(7):641–649, July 1994.

[26] Michael Heinzinger, Ahmed Elnaggar, Yu Wang, Christian Dallago, Dmitrii Nechaev, Florian Matthes, and Burkhard Rost. Modeling aspects of the language of life through transfer-learning protein sequences. BMC Bioinformatics, 20(1):723, December 2019.

[27] Thomas A Hopf, Anna G Green, Benjamin Schubert, Sophia Mersmann, Charlotta P I Schärfe, John B Ingraham, Agnes Toth-Petroczy, Kelly Brock, Adam J Riesselman, Perry Palmedo, Chan Kang, Robert Sheridan, Eli J Draizen, Christian Dallago, Chris Sander, and Debora S Marks. The EVcouplings Python framework for coevolutionary sequence analysis. Bioinformatics, 35(9):1582–1584, May 2019.

[28] Chloe Hsu, Robert Verkuil, Jason Liu, Zeming Lin, Brian Hie, Tom Sercu, Adam Lerer, and Alexander Rives. Learning inverse folding from millions of predicted structures. In Proceedings of the 39th International Conference on Machine Learning, pp. 8946–8970, June 2022.

[29] S.J. Hubbard and J.M. Thornton. NACCESS, Computer Program, 1993.

[30] Zerrin Isik, Christoph Baldow, Carlo Vittorio Cannistraci, and Michael Schroeder. Drug target prioritization by perturbed gene expression and network information. Scientific Reports, 5(1):17417, November 2015.

[31] Justina Jankauskaitė, Brian Jiménez-García, Justas Dapkūnas, Juan Fernández-Recio, and Iain H. Moal. SKEMPI 2.0: an updated benchmark of changes in protein–protein binding energy, kinetics and thermodynamics upon mutation. Bioinformatics, 35(3):462–469, February 2019.

[32] Sherlyn Jemimah, K Yugandhar, and M Michael Gromiha. PROXiMATE: a database of mutant protein–protein complex thermodynamics and kinetics. Bioinformatics, 33(17):2787–2788, September 2017.

[33] H. Jeong, S. P. Mason, A.-L. Barabási, and Z. N. Oltvai. Lethality and centrality in protein networks. Nature, 411(6833):41–42, May 2001.

[34] L. Steven Johnson, Sean R. Eddy, and Elon Portugaly. Hidden Markov model speed heuristic and iterative HMM search procedure. BMC Bioinformatics, 11(1):431, August 2010.

[35] Harry C. Jubb, Arun P. Pandurangan, Meghan A. Turner, Bernardo Ochoa-Montaño, Tom L. Blundell, and David B. Ascher. Mutations at protein-protein interfaces: Small changes over big surfaces have large impacts on human health. Progress in Biophysics and Molecular Biology, 128:3–13, September 2017.

[36] John Jumper, Richard Evans, Alexander Pritzel, Tim Green, Michael Figurnov, Olaf Ronneberger, Kathryn Tunyasuvunakool, Russ Bates, Augustin Žídek, Anna Potapenko, Alex Bridgland, Clemens Meyer, Simon A. A. Kohl, Andrew J. Ballard, Andrew Cowie, Bernardino Romera-Paredes, Stanislav Nikolov, Rishub Jain, Jonas Adler, Trevor Back, Stig Petersen, David Reiman, Ellen Clancy, Michal Zielinski, Martin Steinegger, Michalina Pacholska, Tamas Berghammer, Sebastian Bodenstein, David Silver, Oriol Vinyals, Andrew W. Senior, Koray Kavukcuoglu, Pushmeet Kohli, and Demis Hassabis. Highly accurate protein structure prediction with AlphaFold. Nature, 596(7873):583–589, August 2021.

[37] Elodie Laine and Alessandra Carbone. Local Geometry and Evolutionary Conservation of Protein Surfaces Reveal the Multiple Recognition Patches in Protein-Protein Interactions. PLOS Computational Biology, 11(12):e1004580, 2015.

[38] Elodie Laine, Yasaman Karami, and Alessandra Carbone. GEMME: A Simple and Fast Global Epistatic Model Predicting Mutational Effects. Molecular Biology and Evolution, 36(11):2604–2619, November 2019.

[39] Emmanuel D. Levy. A Simple Definition of Structural Regions in Proteins and Its Use in Analyzing Interface Evolution. Journal of Molecular Biology, 403(4):660–670, November 2010.

[40] Zeming Lin, Halil Akin, Roshan Rao, Brian Hie, Zhongkai Zhu, Wenting Lu, Allan dos Santos Costa, Maryam Fazel-Zarandi, Tom Sercu, Sal Candido, and Alexander Rives. Language models of protein sequences at the scale of evolution enable accurate structure prediction, July 2022. Pages: 2022.07.20.500902 Section: New Results.

[41] Quanya Liu, Peng Chen, Bing Wang, Jun Zhang, and Jinyan Li. dbMPIKT: a database of kinetic and thermodynamic mutant protein interactions. BMC Bioinformatics, 19(1):455, November 2018.

[42] Xianggen Liu, Yunan Luo, Sen Song, and Jian Peng. Pre-training of Graph Neural Network for Modeling Effects of Mutations on Protein-Protein Binding Affinity. arXiv:2008.12473 [cs, q-bio], August 2020.

[43] Céline Marquet, Michael Heinzinger, Tobias Olenyi, Christian Dallago, Kyra Erckert, Michael Bernhofer, Dmitrii Nechaev, and Burkhard Rost. Embeddings from protein language models predict conservation and variant effects. Human Genetics, December 2021.

[44] Joshua Meier, Roshan Rao, Robert Verkuil, Jason Liu, Tom Sercu, and Alex Rives. Language models enable zero-shot prediction of the effects of mutations on protein function. In Advances in Neural Information Processing Systems, volume 34, pp. 29287–29303. Curran Associates, Inc., 2021.

[45] Mihaly Mezei. A new method for mapping macromolecular topography. Journal of Molecular Graphics and Modelling, 21(5):463–472, March 2003.

[46] Milot Mirdita, Konstantin Schütze, Yoshitaka Moriwaki, Lim Heo, Sergey Ovchinnikov, and Martin Steinegger. ColabFold - Making protein folding accessible to all. preprint, doi: 10.21203/rs.3.rs-1032816/v1, November 2021.

[47] Iain H. Moal and Juan Fernández-Recio. SKEMPI: a Structural Kinetic and Energetic database of Mutant Protein Interactions and its use in empirical models. Bioinformatics, 28(20):2600–2607, October 2012.

[48] Yasser Mohseni Behbahani, Simon Crouzet, Elodie Laine, and Alessandra Carbone. Deep Local Analysis evaluates protein docking conformations with locally oriented cubes. Bioinformatics, p. btac551, August 2022.

[49] Saket Navlakha and Carl Kingsford. The power of protein interaction networks for associating genes with diseases. Bioinformatics, 26(8):1057–1063, April 2010.

[50] Surendra S. Negi and Werner Braun. Statistical analysis of physical-chemical properties and prediction of protein-protein interfaces. Journal of Molecular Modeling, 13(11):1157–1167, November 2007.

[51] Hafumi Nishi, Manoj Tyagi, Shaolei Teng, Benjamin A. Shoemaker, Kosuke Hashimoto, Emil Alexov, Stefan Wuchty, and Anna R. Panchenko. Cancer Missense Mutations Alter Binding Properties of Proteins and Their Interaction Networks. PLOS ONE, 8(6):e66273, June 2013.

[52] Guillaume Pagès, Benoit Charmettant, and Sergei Grudinin. Protein model quality assessment using 3D oriented convolutional neural networks. Bioinformatics (Oxford, England), 35(18):3313–3319, September 2019.

[53] Douglas E. V. Pires, David B. Ascher, and Tom L. Blundell. mCSM: predicting the effects of mutations in proteins using graph-based signatures. Bioinformatics (Oxford, England), 30(3):335–342, February 2014.

[54] Douglas E.V. Pires and David B. Ascher. mCSM-AB: a web server for predicting antibody–antigen affinity changes upon mutation with graph-based signatures. Nucleic Acids Research, 44(W1):W469–W473, July 2016.

[55] Janet Piñero, Núria Queralt-Rosinach, Àlex Bravo, Jordi Deu-Pons, Anna Bauer-Mehren, Martin Baron, Ferran Sanz, and Laura I. Furlong. DisGeNET: a discovery platform for the dynamical exploration of human diseases and their genes. Database, 2015(bav028), January 2015.

[56] Roshan Rao, Joshua Meier, Tom Sercu, Sergey Ovchinnikov, and Alexander Rives. Transformer protein language models are unsupervised structure learners. bioRxiv, p. 2020.12.15.422761, December 2020.

[57] Raffaele Raucci, Elodie Laine, and Alessandra Carbone. Local Interaction Signal Analysis Predicts Protein-Protein Binding Affinity. Structure, 26(6):905–915.e4, June 2018.

[58] Alexander Rives, Joshua Meier, Tom Sercu, Siddharth Goyal, Zeming Lin, Jason Liu, Demi Guo, Myle Ott, C. Lawrence Zitnick, Jerry Ma, and Rob Fergus. Biological structure and function emerge from scaling unsupervised learning to 250 million protein sequences. Proceedings of the National Academy of Sciences, 118(15), April 2021.

[59] Carlos H. M. Rodrigues, Yoochan Myung, Douglas E. V. Pires, and David B. Ascher. mCSM-PPI2: predicting the effects of mutations on protein–protein interactions. Nucleic Acids Research, 47(W1):W338–W344, July 2019.

[60] Carlos H M Rodrigues, Douglas E V Pires, and David B Ascher. mmCSM-PPI: predicting the effects of multiple point mutations on protein–protein interactions. Nucleic Acids Research, 49(W1):W417–W424, July 2021.

[61] Maxim V. Shapovalov and Roland L. Dunbrack. A Smoothed Backbone-Dependent Rotamer Library for Proteins Derived from Adaptive Kernel Density Estimates and Regressions. Structure, 19(6):844–858, June 2011.

[62] Rohit Singh, Kapil Devkota, Samuel Sledzieski, Bonnie Berger, and Lenore Cowen. Topsy-Turvy: integrating a global view into sequence-based PPI prediction. Bioinformatics, 38(Supplement_1):i264–i272, July 2022.

[63] Sarah Sirin, James R. Apgar, Eric M. Bennett, and Amy E. Keating. AB-Bind: Antibody binding mutational database for computational affinity predictions. Protein Science, 25(2):393–409, 2016.

[64] Colin A. Smith and Tanja Kortemme. Backrub-Like Backbone Simulation Recapitulates Natural Protein Conformational Variability and Improves Mutant Side-Chain Prediction. Journal of Molecular Biology, 380(4):742–756, July 2008.

[65] Kai Sun, Joana P. Gonçalves, Chris Larminie, and Nataša Pržulj. Predicting disease associations via biological network analysis. BMC Bioinformatics, 15(1):304, September 2014.

[66] Baris E. Suzek, Yuqi Wang, Hongzhan Huang, Peter B. McGarvey, Cathy H. Wu, and the UniProt Consortium. UniRef clusters: a comprehensive and scalable alternative for improving sequence similarity searches. Bioinformatics, 31(6):926–932, March 2015.

[67] Xiwei Tang, Qiu Xiao, and Kai Yu. Breast Cancer Candidate Gene Detection Through Integration of Subcellular Localization Data With Protein–Protein Interaction Networks. IEEE Transactions on NanoBioscience, 19(3):556–561, July 2020.

[68] Shaolei Teng, Thomas Madej, Anna Panchenko, and Emil Alexov. Modeling Effects of Human Single Nucleotide Polymorphisms on Protein-Protein Interactions. Biophysical Journal, 96(6):2178–2188, March 2009.

[69] G. C. P. van Zundert, J. P. G. L. M. Rodrigues, M. Trellet, C. Schmitz, P. L. Kastritis, E. Karaca, A. S. J. Melquiond, M. van Dijk, S. J. de Vries, and A. M. J. J. Bonvin. The HADDOCK2.2 Web Server: User-Friendly Integrative Modeling of Biomolecular Complexes. Journal of Molecular Biology, 428(4):720–725, February 2016.

[70] Anna Vangone and Alexandre MJJ Bonvin. Contacts-based prediction of binding affinity in protein–protein complexes. eLife, 4:e07454, July 2015.

[71] Ashish Vaswani, Noam Shazeer, Niki Parmar, Jakob Uszkoreit, Llion Jones, Aidan N Gomez, Łukasz Kaiser, and Illia Polosukhin. Attention is All you Need. In Advances in Neural Information Processing Systems, volume 30. Curran Associates, Inc., 2017.

[72] Thom Vreven, Iain H. Moal, Anna Vangone, Brian G. Pierce, Panagiotis L. Kastritis, Mieczyslaw Torchala, Raphael Chaleil, Brian Jiménez-García, Paul A. Bates, Juan Fernandez-Recio, Alexandre M. J. J. Bonvin, and Zhiping Weng. Updates to the Integrated Protein–Protein Interaction Benchmarks: Docking Benchmark Version 5 and Affinity Benchmark Version 2. Journal of Molecular Biology, 427(19):3031–3041, September 2015.

[73] Menglun Wang, Zixuan Cang, and Guo-Wei Wei. A topology-based network tree for the prediction of protein–protein binding affinity changes following mutation. Nature Machine Intelligence, 2(2):116–123, February 2020. Number: 2 Publisher: Nature Publishing Group.

[74] Dapeng Xiong, Dongjin Lee, Le Li, Qiuye Zhao, and Haiyuan Yu. Implications of disease-related mutations at protein–protein interfaces. Current Opinion in Structural Biology, 72:219–225, February 2022.

[75] Peng Xiong, Chengxin Zhang, Wei Zheng, and Yang Zhang. BindProfX: Assessing Mutation-Induced Binding Affinity Change by Protein Interface Profiles with Pseudo-Counts. Journal of Molecular Biology, 429(3):426–434, February 2017.

[76] Ning Zhang, Yuting Chen, Haoyu Lu, Feiyang Zhao, Roberto Vera Alvarez, Alexander Goncearenco, Anna R. Panchenko, and Minghui Li. MutaBind2: Predicting the Impacts of Single and Multiple Mutations on Protein-Protein Interactions. iScience, 23(3):100939, March 2020.

[77] Zuobai Zhang, Minghao Xu, Arian Jamasb, Vijil Chenthamarakshan, Aurelie Lozano, Payel Das, and Jian Tang. Protein Representation Learning by Geometric Structure Pretraining, May 2022. arXiv:2203.06125 [cs].

[78] Guangyu Zhou, Muhao Chen, Chelsea J T Ju, Zheng Wang, Jyun-Yu Jiang, and Wei Wang. Mutation effect estimation on protein–protein interactions using deep contextualized representation learning. NAR Genomics and Bioinformatics, 2(2):lqaa015, June 2020.

